# Molecular basis of the biogenesis of a protein organelle for ethanolamine utilization

**DOI:** 10.1101/2024.05.17.594633

**Authors:** Mengru Yang, Oluwatobi Adegbite, Ping Chang, Xiaojun Zhu, Yan Li, Gregory F. Dykes, Yu Chen, Natasha Savage, Jay C. D. Hinton, Lu-Yun Lian, Lu-Ning Liu

## Abstract

Many pathogenic bacteria use proteinaceous ethanolamine-utilization microcompartments (Eut BMCs) to facilitate the catabolism of ethanolamine, an abundant nutrient in the mammalian gut. The ability to metabolize ethanolamine gives pathogens a competitive edge over commensal microbiota which can drive virulence in the inflamed gut. Despite their critical functions, the molecular mechanisms underlying the synthesis of Eut BMCs in bacterial cells remain elusive. Here, we report a systematic study for dissecting the molecular basis underlying Eut BMC assembly in *Salmonella*. We determined the functions of individual building proteins in the structure and function of Eut BMCs and demonstrated that EutQ plays an essential role in both cargo encapsulation and Eut BMC formation through specific association with the shell and cargo enzymes. Furthermore, our data reveal that Eut proteins can self-assemble to form cargo and shell aggregates independently *in vivo*, and that the biogenesis of Eut BMCs follows a unique “Shell-first” pathway. Cargo enzymes exhibit dynamic liquid-like organization within the Eut BMC. These discoveries provide mechanistic insights into the structure and assembly of the Eut BMC, which serves as a paradigm for membrane-less organelles. It opens up new possibilities for therapeutic interventions for infectious diseases.

## INTRODUCTION

Ethanolamine (EA) is an abundant nutrient in the membrane-rich gastrointestinal (GI) tract, attributed to the constant turnover of the mucosal epithelium and the resident microbiota ^1-3^. Diverse gut bacteria, including those from pathogenic genera such as *Salmonella, Clostridium, Klebsiella, Listeria, Escherichia*, and *Enterococcus*, can utilize EA as a sole source of carbon and/or nitrogen ^4-6^. The ability to metabolize EA provides a competitive growth advantage to pathogens over commensal microbiota. Moreover, EA plays a role in regulating the virulence of these pathogens ^7,8^ and EA catabolism is closely associated with virulence in the inflamed gut ^9-13^.

In many pathogenic species, such as *Salmonella enterica* serovar Typhimurium (*S*. Typhimurium), EA utilization is executed by a group of proteins that are encoded by 17 genes clustered in an *eut* operon ^14-16^. These proteins, known as EutS/P/Q/T/D/M/N/E/J/G/H/A/B/C/L/K/R, include catalytic and structural protein components that self-assemble to form a polyhedral organelle, called the ethanolamine-utilization microcompartment (Eut BMCs, Figs. 1A and 1B). The Eut BMCs belong to a family of bacterial microcompartments (BMCs) that are widespread in the bacterial kingdom, which perform a variety of functions, including CO_2_ fixation, infection, and microbial ecology ^16-21^. However, despite their important functions in bacterial pathogenesis and host–pathogen interactions, our understanding of how the various shell proteins and cargo enzymes interact and self-assemble *in vivo* to form functional Eut BMCs is still limited. Here, we perform a systematic study on the molecular mechanisms that underlie the organization and biogenesis of Eut BMCs in *S*. Typhimurium LT2 strain. Using genetic modification, super-resolution fluorescence imaging, electron microscopy (EM), and growth assays, we delineate the roles of individual Eut components in Eut BMC assembly and functionality. Combining NMR spectroscopy, AlphaFold predictions^22^, and biochemical studies, we elucidate how the N-terminal domain of EutQ mediates the physical binding between the shell and cargos to ensure encapsulation. We further report that Eut proteins independently self-assemble to form cargo and shell aggregates. Our results establish that the *in vivo* biogenesis of Eut BMCs follows a distinct “Shell first” assembly pathway, in which the assembly of shells take place before the assembly of cargos and at the cell poles. We also demonstrate that the Eut BMC cargo core possesses liquid-like features, highlighting the significance of phase separation in constructing and maintaining self-assembling protein organelles in pathogenic bacteria. Our study addresses the long-standing questions about how Eut BMCs are generated and how the shell and cargos self-assemble to form functional protein organelles. The new knowledge could guide the development of therapeutic interventions for diseases caused by pathogens.

**Fig. 1.**
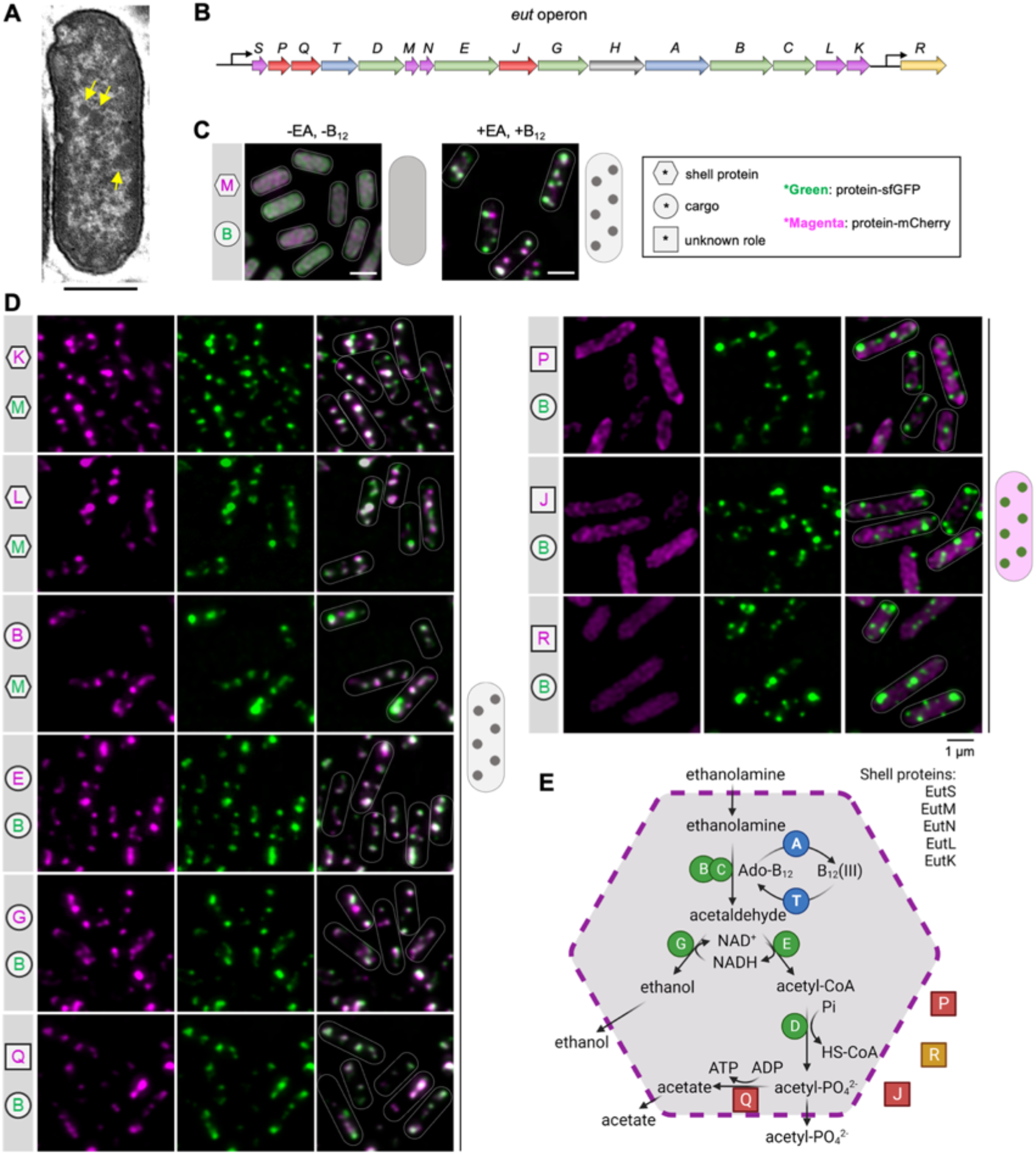
Eut BMC structure, gene operon, and protein organization in *S*. Typhimurium LT2. **A**, Thin-section transmission electron microscopy (TEM) of the *S*. Typhimurium LT2 wild-type (WT) strain following growth in the presence of EA and B_12_, showing the formation of Eut BMCs (yellow arrows). Scale bar: 500 nm. **B**, The chromosomal *eut* operon of *S*. Typhimurium LT2 includes structural genes (*eutSMNLK*), catalytic genes (*eutDEGBC* for EA degradation, and the *eutTA* for B_12_ recycling), *eutR* (transcriptional regulator), and unknown function genes (*eutPQJ*), which are indicated in distinct colors as show in panel **E. C**, Fluorescence images show the *S*. Typhimurium LT2 cells carrying pBAD-*eutM-mCherry*::*eutB-sfGFP* grown in minimal medium, in the absence or presence of EA and B_12_. Hexagon represents shell proteins; circle represents cargo; square represents proteins with an unknown role and EutR. Eut protein in green represents tagging with sfGFP, while that in magenta represents tagging with mCherry. Grey color represents that the mCherry and sfGFP signals are colocalized. Scale bar: 1 µm. **D**, Localization of Eut proteins in the cells grown in minimal medium in the presence of EA and B_12_. **E**, Schematic model of the protein organization of the Eut BMC (created with BioRender). The color scheme is the same as shown in panel **B**, indicating the different organization of individual Eut proteins.

## RESULTS

### Subcellular localization of Eut proteins and protein organization of the Eut BMC

To examine the subcellular location and assembly of Eut proteins in *S*. Typhimurium, we labeled target Eut proteins with fluorescent proteins using a pBAD/*Myc*-His vector with the arabinose-inducible P*araBAD* promoter ^23^ (Fig. 1C, Fig. S1, Tables S1 and S2). For *in vivo* super-resolution fluorescence imaging, selected Eut shell or cargo proteins tagged with superfolder green fluorescent protein (sfGFP) or mCherry were expressed under the control of the P*araBAD* promoter, along with the expression of the endogenous *eut* operon, which was activated in the presence of EA and cobalamin (vitamin B_12_) ^24^.

In the absence of EA and B_12_, no endogenous Eut proteins were expressed in *S*. Typhimurium. In the absence of Eut BMC formation, the individual vector-expressed Eut proteins, including the shell proteins EutM/K/L, cargo proteins EutG/B, as well as EutQ/P/J and EutR, were localized throughout the cytoplasm (Fig. 1C, Fig. S2). One exception is the cargo protein EutE, which formed an aggregate at the cell pole. In addition, the multiple enzyme complex EutBC (sfGFP was fused at the EutC C-terminus) appeared to aggregate at a single pole of the cell (Fig. S3). Both EutE and EutC possess short peptides at their N-termini, which act as endogenous encapsulation peptides (EPs) to target cargo enzymes into the Eut BMC ^25,26^. The observed polar aggregation of EutE and EutBC was likely mediated by their EPs, consistent with the findings from 1,2-propanediol-utilization microcompartments (Pdu BMCs) ^23^.

Growth of *S*. Typhimurium in the presence of EA and B_12_ led to expression of endogenous Eut proteins and formation of Eut BMCs (Fig. 1A). Fluorescence imaging revealed that the Eut proteins (shell proteins EutM/K/L and cargo proteins EutE/G/B/Q) exhibited the patchy distribution, representing the signature distribution of Eut BMCs *in vivo* (Figs. 1C, 1D). By contrast, EutP, EutJ, and EutR were evenly distributed throughout the cytoplasm, indicating that these proteins are not incorporated within Eut BMCs. EutQ and EutP were assumed to function as acetate kinases ^27^. Our results show that only EutQ is encapsulated within the Eut BMC, possibly catalyzing the formation of acetate and ATP from acetyl phosphate generated from EutD (see further analysis below). Our data lead us to propose a model for the protein organization of the Eut BMC (Fig. 1E).

### Roles of individual Eut proteins in Eut BMC assembly

To decipher the roles of individual protein components in the Eut BMC assembly, we generated a series of mutants that lacked individual Eut proteins via scarless deletion of every *eut* gene (Δ*eutS*, Δ*eutM*, Δ*eutN*, Δ*eutL*, Δ*eutK*, Δ*eutQ*) (Figs. S4, S5A). The dual-labeling plasmid pBAD-*eutM-mCherry*::*eutB-sfGFP* was then transformed into these deletion mutants to visualize the assembly of Eut shell proteins (indicated by EutM) and cargos (indicated by EutB).

In the absence of EutL or EutQ, EutB-sfGFP aggregated into a large punctum at a cell pole, whereas EutM-mCherry formed multiple clustered assemblies in the cytoplasm, suggesting spatial separation of shell and cargo proteins (Figs. 2A, 2B, Fig. S6). We conclude that both EutL and EutQ play essential roles in mediating association between the shell and cargo proteins of Eut BMCs. Because the separation of shell and cargo proteins leads to the release of toxic acetaldehyde, the growth rates of the Δ*eutL* and Δ*eutQ* strains were reduced compared with the wild-type (WT) strain (Fig. 2G, Fig. S7). The Δ*eutQ* grew even slower than Δ*eutL*, consistent with EutQ acting as an acetate kinase during EA degradation ^27^. Expression of EutL or EutQ in the Δ*eutL* or Δ*eutQ* strain, respectively, partially rescued the assembly of Eut BMCs, evidenced by colocalization of EutM-mCherry and EutB-sfGFP and growth assays (Fig. S8). We found that complementation of EutL resulted in various aggregates mostly at cell poles, whereas complementation of EutQ led to formation of rod-like structures (Fig. S8A). These results suggest that precise levels of EutL/EutQ are required to define the assembly of native Eut BMCs.

**Fig. 2.**
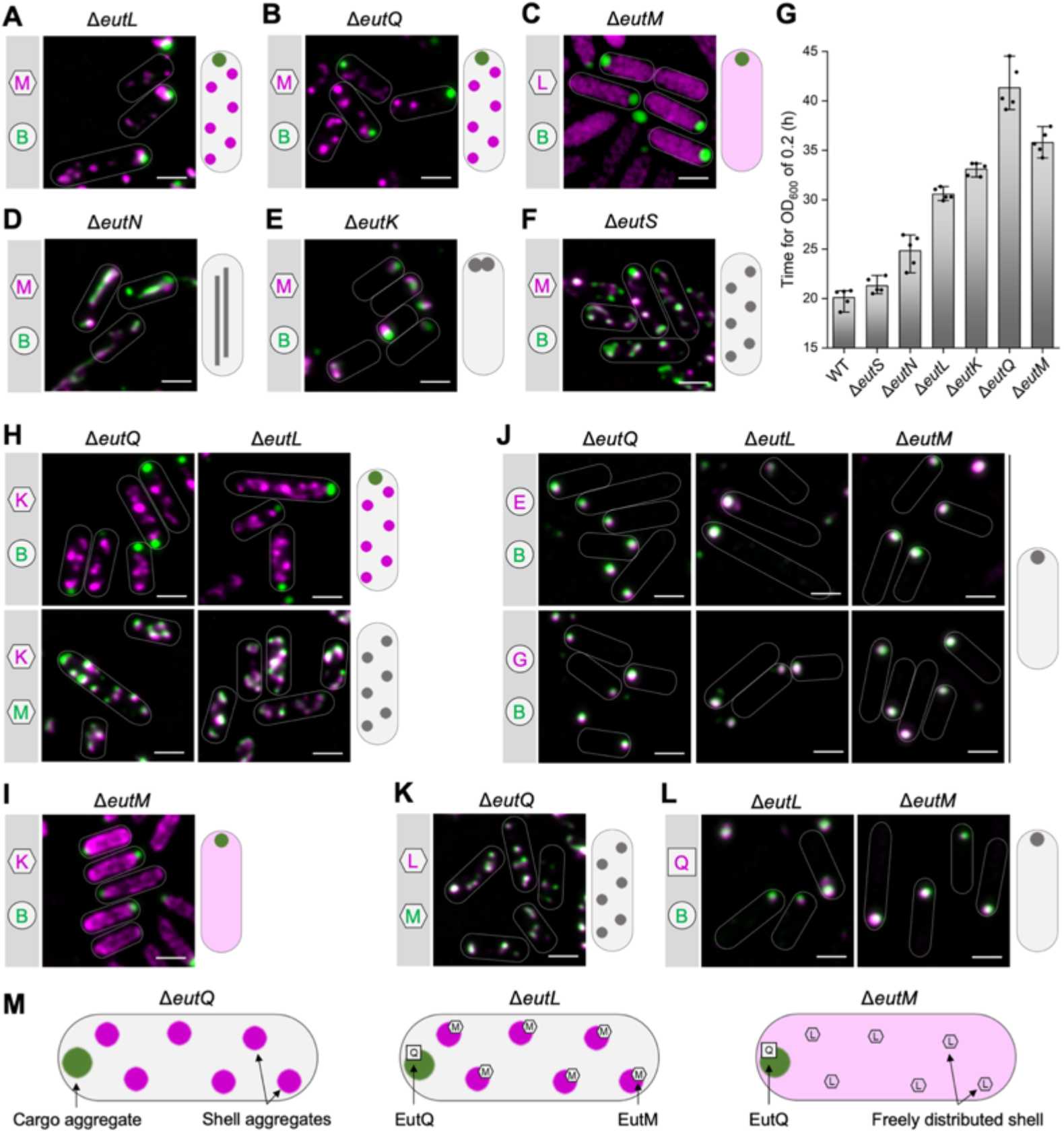
Roles of *eut* operon-encoding proteins in the assembly of Eut BMCs in *S*. Typhimurium LT2. **A-F**, Localization of shell proteins and cargos in the gene deletion mutants in the absence of EutL, EutQ, EutM, EutN, EutK, or EutS, which were grown in the presence of EA and B_12_. EutM-mCherry (shell) and EutB-sfGFP (cargo) were imaged to indicate the locations of shell proteins and cargos. In the Δ*eutM* mutant, EutL-mCherry and EutB-sfGFP were imaged to indicate the locations of shell proteins and cargos. Schematic models of the *in vivo* localization of shell and cargo assemblies are shown on the right. **G**, Time for LT2-WT and mutants to grow to reach OD_600_=0.2 in M9 medium supplemented with EA and B_12_ (*n*=5). The center for error bars represents the mean. The whiskers extend to the smallest and largest data points that are within 1.5 times the interquartile range of the upper and lower quartiles. *n* number of biologically independent experiments. **H-J**, Locations of shell and cargo proteins in the Δ*eutQ*, Δ*eutL* and Δ*eutM* strains. **K**, Distribution of EutL-mCherry and EutM-sfGFP in Δ*eutQ*. **L**, Distribution of EutQ-mCherry and EutB-sfGFP in Δ*eutL* and Δ*eutM*. **M**, Models for the locations of Eut proteins in Δ*eutQ*, Δ*eutL* and Δ*eutM*. All strains were imaged following growth in the minimum medium in the presence of EA and B_12_. Scale bars: 1 µm.

For the Δ*eutM* strain, we used the pBAD-*eutL-mCherry*::*eutB-sfGFP* to monitor the assembly of shell proteins (represented by EutL) and cargos (represented by EutB) (Fig. 2C). In the absence of EutM, EutB-sfGFP formed a large punctum at a single polar position within the cell, whereas EutL-mCherry was evenly distributed throughout the cytosol (Fig. 2C, Fig. S6). The Δ*eutM* strain exhibited a growth defect on EA compared to the WT (Fig. 2G). These results indicate that EutM may initiate the assembly of the Eut BMC shell.

In the Δ*eutN* mutant, Eut assemblies were elongated (Fig. 2D, Fig. S6), suggesting that EutN proteins play an important role in shaping the Eut BMC structure. This aligns with the functions of shell vertex pentameric proteins in other BMCs, including CsoS4 in the α-carboxysome ^28^, CcmL in the β-carboxysome ^29^, and PduN in the Pdu BMC ^23,30^. EutN is the only CsoS4/CcmL homolog in *Salmonella enterica*, and no other pentamers have been identified in Eut BMCs ^31^. Interestingly, EutN was characterized to be pentameric in solution ^32^, whereas crystallization analysis showed that EutN was a homohexamer ^29,33^, suggesting the quaternary structural flexibility of the shell proteins under varying environments.

In Δ*eutK*, Eut BMCs were predominantly located at cell poles (Fig. 2E, Fig. S6) and these polar Eut BMC assemblies were more static than the highly dynamic Eut BMCs in the WT (Supplemental Videos 1 and 2), indicating that EutK plays a role in determining the spatial distribution of Eut BMCs in *S*. Typhimurium. The Δ*eutK* strain exhibited a slower growth rate than the WT on EA (Fig. 2G, Fig. S7). Complementation experiments showed that Eut BMC formation and cell growth recovered when EutK was expressed in Δ*eutK* (Fig. S8).

In the absence of EutS, EutM-mCherry and EutB-sfGFP colocalized into fluorescent foci (Fig. 2F), resembling the typical Eut BMC distribution in WT cells (Fig. 1C). Thin-section EM confirmed the formation of Eut BMC in Δ*eutS* (Fig. S6). The Δ*eutS* strain exhibited a similar growth profile as the WT using EA as sole carbon source (Fig. 2G, Fig. S7), consistent with previous observations ^34^. These results suggest that EutS is not essential for the assembly of Eut BMCs.

To sum up, our systematic analysis revealed the essential roles of EutM, EutN, EutL, EutK, and EutQ for determining the structural integrity and growth of Eut BMCs.

### EutQ governs the encapsulation of the enzymatic core to the shell

Our results showed that in the absence of EutL, EutQ, and EutM, the cargo enzymes were spatially separated from the shell proteins; cargos formed one aggregate at the cell pole, whereas shell proteins appeared as multiple aggregates (Δ*eutL* and Δ*eutQ*) or were evenly distributed (Δ*eutM*) in cells (Fig. 2A-C). To investigate the protein composition of these self-assemblies, we determined the locations of Eut proteins in the Δ*eutQ*, Δ*eutL*,and Δ*eutM* strains grown upon EA and B_12_.

In Δ*eutQ* and Δ*eutL*, the shell protein EutK was dissociated from the cargo EutB, but colocalized with the shell protein EutM to form multiple aggregates (Fig. 2H). In Δ*eutM*, EutK was freely distributed in the cytoplasm (Fig. 2I), which fits with the observations for EutL (Fig. 2C), confirming that EutM drives shell formation. Furthermore, in Δ*eutQ*, Δ*eutL*, and Δ*eutM*, the cargo proteins EutE and EutG colocalized with EutB, forming a single polar aggregate (Fig. 2J). These findings indicate that the lack of EutQ, EutL, or EutM could cause shell-cargo disassociation, resulting in the formation of a cargo aggregate at the cell pole coupled with multiple shell aggregates or the dispersion of shell proteins within cells (Figs. 2A-C, 2H-J, 2M).

Our results suggest that interactions between EutQ and EutL or/and EutM may be essential for Eut BMC cargo-shell association. To test this hypothesis, we examined the locations of EutL and EutM in Δ*eutQ*, and the distribution of EutQ in Δ*eutL* and Δ*eutM*. In the absence of EutQ (Δ*eutQ*), EutL and EutM assembled together to form shell aggregates (Fig. 2K) but disassociated with cargos (Figs. 2B, 2H). In Δ*eutL* and Δ*eutM*, EutQ formed a punctum at the cell pole that colocalized with cargos (Fig. 2L). We conclude that EutQ forms a strong association with cargos while interacting with the shell proteins EutM and EutL, which is essential for cargo encapsulation. The findings from Δ*eutL* show that shell assemblies disassociated with cargos in the presence of EutM (Figs. 2H, 2M), suggesting that the interactions between EutQ and EutM were insufficient for cargo encapsulation. This differs from previous pull-down assays, which suggested that the N-terminus of EutQ could only interact with EutM among all shell proteins ^35^. In Δ*eutM*, EutL displayed the cytoplasmic distribution, implying that without EutM, EutL does not bind strongly with EutQ (Fig. 2C, 2M). We propose that the binding of EutQ to the shell is facilitated by the shell proteins EutL and EutM, and that EutQ specifically attaches at the interaction interface between EutM and EutL.

### Molecular mechanisms of the EutQ-mediated shell-cargo association

Size exclusion chromatography coupled with multi-angle light scattering analysis revealed that EutQ (∼25 KDa per monomer) forms a ∼50 KDa dimer, and that dimerization was mediated by the contacts between two EutQ C-termini (Fig. S9). This is consistent with the crystal structure of EutQ C-terminus from *Clostridioides difficile* 630 (PDB ID: 2PYT) ^36^. Secondary structure prediction using Phyre2 ^37^ indicated that the EutQ N-terminal region (EutQ^1-99^) possesses four α-helices (H1-H4) and a long flexible unstructured region (41-61 amino acid residues) linking H2 and H3 (Fig. 3A). AlphaFold prediction of the entire EutQ structure shows a similar 4-α-helix linked by a long flexible segment for EutQ^1-99^, followed by the C-terminus region (EutQ^98-229^) that consists of mainly β-strands (Figs. S10A and B).

**Fig. 3.**
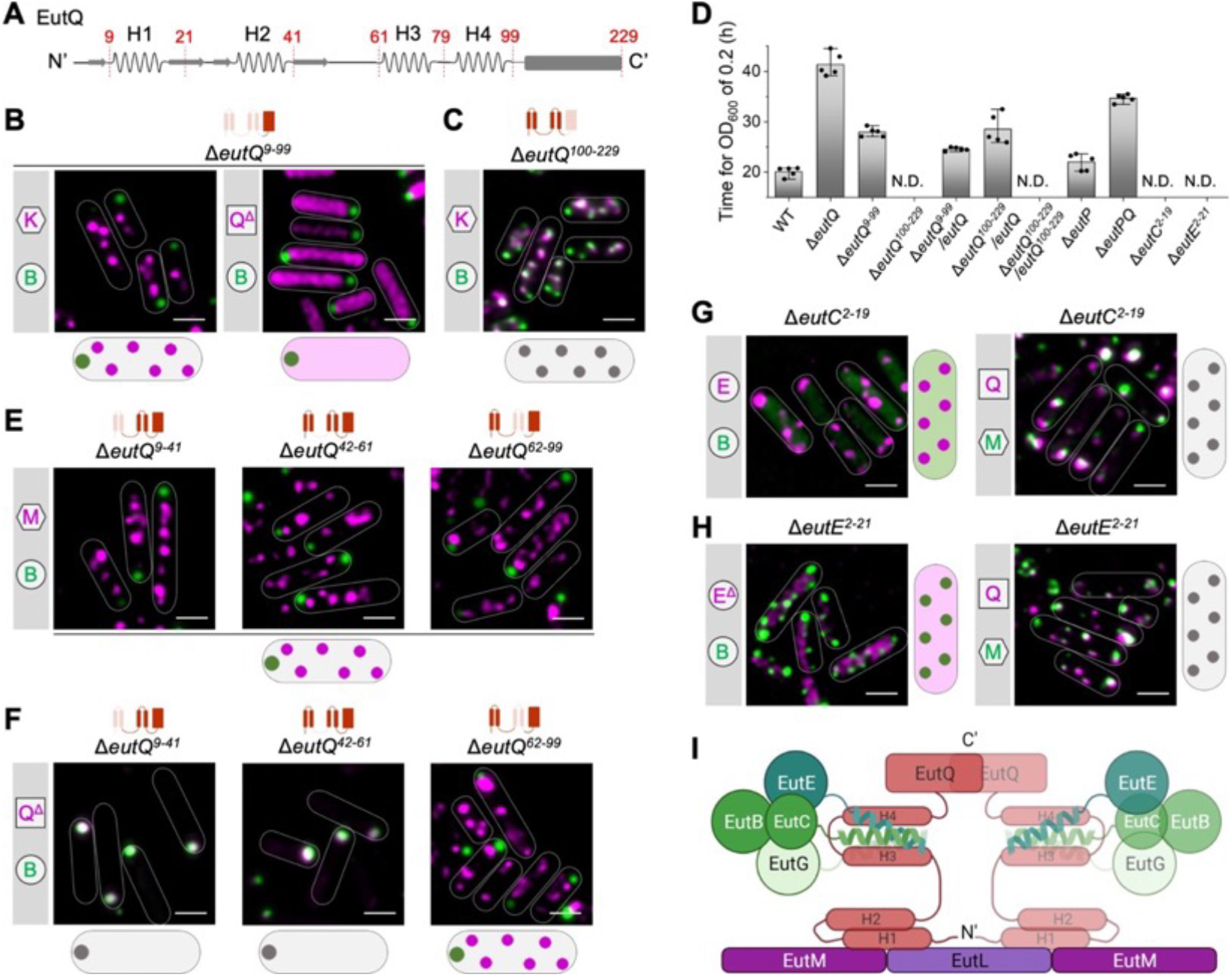
EutQ plays an essential role in mediating the binding of the shell and cargos. **A**, Secondary structure of EutQ with the N-terminus predicted by Phyre2 ^37^. H1-H4 represent four α-helix structures. The amino acid sites for genome editing were labeled and colored in red. **B**, EutK-mCherry (shell) and EutB-sfGFP (cargo), and EutQ^Δ^-mCherry and EutB-sfGFP (cargo) were visualized in Δ*eutQ*^*9-99*^ in the presence of EA and B_12_. Schematic models of the *in vivo* localization of shell and cargo assemblies are shown at the bottom. **C**, EutK-mCherry (shell) and EutB-sfGFP (cargo) were visualized in Δ*eutQ*^*100-229*^ in the presence of EA and B_12_. **D**, Time for LT2-WT and mutants to grow to OD_600_=0.2 on EA and B_12_ in M9 medium (*n*=5). The center for error bars represents the mean. The whiskers extend to the smallest and largest data points that are within 1.5 times the interquartile range of the upper and lower quartiles. *n* number of biologically independent experiments. **E**, EutM-mCherry (shell) and EutB-sfGFP (cargo) were visualized in Δ*eutQ*^*9-41*^ Δ*eutQ*^*42-61*^, and Δ*eutQ*^*62-99*^ in the presence of EA and B_12_. **F**, EutQ^Δ^-mCherry and EutB-sfGFP (cargo) were visualized in Δ*eutQ*^*9-41*^, Δ*eutQ*^*42-61*^, and Δ*eutQ*^*62-99*^ in the presence of EA and B_12_. EutQ^Δ^ represents the mutated EutQ with different fragments deleted, which differs from each other in different genome editing mutants. The deletion parts of EutQ were shown in the models on the top of **B, C, E** and **F. G**, EutE-mCherry/EutB-sfGFP and EutQ-mCherry/EutM-sfGFP were visualized in Δ*eutC*^*2-19*^ following growth in minimal medium in the presence of EA and B_12_. **H**, EutE^Δ^-mCherry/EutB-sfGFP and EutQ-mCherry/EutM-sfGFP were visualized in Δ*eutE*^*2-21*^ following growth in minimal medium in the presence of EA and B_12_. EutE^Δ^ represents the C-terminus of EutE with EutE^2-21^ deleted. **I**, Schematic model for the assembly mode between EutQ and shell/cargo (created with BioRender). EutQ is shown as a dimer, and the dimerization site is at the C-terminus. Scale bar in fluorescence images: 1 µm.

To determine the function of EutQ in Eut BMC assembly, we generated two mutants that lacked the EutQ N-terminus (Δ*eutQ*^*9-99*^, note that *eutQ*^*1-9*^ overlaps with the upstream gene *eutP*) or the C-terminus (Δ*eutQ*^*100-229*^). Deletion of EutQ^9-99^ caused disassociation of the shell and cargos and formation of a polar cargo aggregate along with numerous shell assemblies (Fig. 3B, Figs. S10C, S10E), in line with the observations from the deletion of full-length EutQ (Fig. 2, Fig. S6). In the absence of EutQ^9-99^ (Δ*eutQ*^*9-99*^), the EutQ C-terminus exhibited an even distribution throughout the cytosol, and did not assemble with either the shells or the cargos (Fig. 3B). By contrast, Eut BMCs were formed in Δ*eutQ*^*100-229*^ (Fig. 3C, Figs. S10D, S10F). These results demonstrate that the EutQ N-terminus plays a critical role in mediating the shell-cargo association, whereas its C-terminus is not essential for Eut BMC assembly.

Both the Δ*eutQ*^*9-*99^ and Δ*eutQ* mutants had reduced growth rates compared to the WT (Fig. 3D, Fig. S11). We propose that the disassociation of cargos and the shell observed in Δ*eutQ*^*9-99*^ and Δ*eutQ* results in accumulation of the toxic metabolic intermediate acetaldehyde. Both EutQ and EutP have been suggested to have acetate kinase activity, facilitating the conversion of acetate and ATP into acetyl-phosphate and vice versa, and EutQ, but not EutP, is required for *Salmonella* growth ^27^. Likewise, EutQ was found to be 10 times more abundant than EutP in *Escherichia coli* ^38^. Our results demonstrate that EutQ is incorporated within the Eut BMC, whereas EutP is distributed in the cytosol (Fig. 1D). We found that Δ*eutP* had a similar growth rate to the WT, and surprisingly that Δ*eutPQ* showed slightly faster growth than Δ*eutQ* (Fig. S11). This presumably reflects the presence of the housekeeping acetate kinase (AckA) in the cytosol, which is encoded by the *ackA* gene out with the *eut* operon ^39,40^. We propose that EutQ can bind with acetyl-phosphate within the Eut BMC, generating a higher catalytic efficiency than EutP and AckA. In Δ*eutQ*^*9-99*^, formation of acetate can be catalyzed by EutQ^100-229^ that contains the C-terminal catalytic domain of EutQ, ensuring a faster growth of Δ*eutQ*^*9-*99^ than Δ*eutQ* (Fig. 3D, Fig. S11). Moreover, the absence of EutQ^100-229^ ablated cell growth for over 50 hours when EA was the sole carbon source. In Δ*eutQ*^*100-229*^, Eut BMCs were still formed. However, the access of acetyl-phosphate to the external acetate kinase was limited by shell encapsulation, resulting in a slower growth rate of Δ*eutQ*^*100-229*^ compared to other strains (Fig. 3D, Fig. S11). The growth of Δ*eutQ*^*100-229*^ was restored by expressing EutQ (Fig. 3D, Fig. S11). However, expression EutQ^100-229^ in Δ*eutQ*^*100-229*^ did not restore growth, suggesting that EutQ^100-229^ itself without the EutQ N-terminus could not be encapsulated into the Eut BMC.

To determine how the N-terminus of EutQ mediates shell-cargo association, we deleted the H1-H4 α-helices individually (Δ*eutQ*^*9-21*^, Δ*eutQ*^*22-41*^, Δ*eutQ*^*62-79*^, and Δ*eutQ*^*80-99*^) and examined the subcellular locations of shell and cargo proteins in these mutants. Deletion of each α-helix led to the disassociation of the shell and cargos (Fig. S12), indicating that all of four α-helices are essential for Eut BMC formation. The AlphaFold-predicted EutQ structure and EutQ^1-99^-EutC^1-20^ interactions (Figs. S10A, S13) led us to propose that H1/H2 interacts with the shell and that H3/H4 is responsible for anchoring the EPs of cargos. To test this hypothesis, we deleted the H1/H2 (Δ*eutQ*^*9-41*^), H3/H4 (Δ*eutQ*^*62-99*^), and the flexible region (Δ*eutQ*^*42-61*^), individually. The shell and cargo assemblies were spatially separated in all three mutants (Fig. 3E, Fig. S14), suggesting that the three regions were crucial for shell-cargo association. Furthermore, EutQ^Δ9-41^ and EutQ^Δ42-61^ aggregated with the cargo cores at the cell poles in Δ*eutQ*^*9-41*^and Δ*eutQ*^*42-61*^, whereas EutQ^Δ62-99^ dissociated with the cargos and colocalized with the shell assemblies in Δ*eutQ*^*62-99*^ (Fig. 3F, Fig. S15).

Together, our results demonstrate that the EutQ N-terminal H1/H2 interacts with the shell while H3/H4 associates with cargos. We propose that the interactions between H3/H4 and cargos were stronger than those between H1/H2 and the shell, because the EutQ depleted of the flexible region (EutQ^Δ42-61^) colocalized with cargos rather than shells (Fig. 3F, middle). This finding indicates that the flexible region plays an important role in shell-cargo binding and potential interactions with the shell.

We then examined the roles of endogenous cargo-derived EPs by studying their interactions with EutQ. Deletion of the N-terminus of EutC (Δ*eutC*^*2-19*^) caused cytosolic distribution of EutB, whereas EutQ, the cargo protein EutE, and the shell protein EutM could still form assemblies, individually (Fig. 3G). These results suggest that the EutC N-terminus was specifically responsible for the assembly and encapsulation of EutBC cargo complexes. Likewise, removal of the native EP of EutE (Δ*eutE*^*2-21*^) resulted in even distribution of EutE^Δ2-21^ throughout the cytosol, while the subcellular locations of other Eut proteins were not affected (Fig. 3H), identifying a role for EutE^2-21^ in targeting EutE to the Eut BMC. Growth assays of the EP-deletion mutants showed no growth on EA as sole carbon source after 50 hours (Fig. 3D, Fig. S11), indicating that without the EPs, EutBC and EutE could not be incorporated into Eut BMCs. The location of another cargo protein, EutG, was not affected in both Δ*eutC*^*2-19*^ and Δ*eutE*^*2-21*^, indicating that EutG possesses its own EP (EutG^1-20^) for recruitment into the Eut BMC (Fig. S16). Collectively, our results unravel the mechanisms by which the EutQ N-terminus mediates the assembly of the Eut shell and cargos (Fig. 3I).

To determine how EutQ binds cargo EPs, we examined the protein-protein interactions using isothermal titration calorimetry (ITC) and nuclear magnetic resonance (NMR) spectroscopy. The ITC results revealed strong binding between EutC^1-20^ and EutQ (*K*_d_ = 5.50 μM), or between EutC^1-20^ and the N-terminal region of EutQ, EutQ^1-99^, fused with a soluble fusion partner, the immunoglobulin binding domain of streptococcal protein G (GB1) ^41^ (*K*_d_ = 7.60 μM) (Figs. 4A, 4B, Fig. S17). A binding stoichiometry of 2 for the EutQ interactions suggests that EutQ functions as a dimer. This is in contrast to EutQ^1-99^-GB1, where the binding stoichiometry to the same EutC peptide is 1, consistent with the monomeric state of EutQ^1-99^. No binding was detected between EutC^1-20^ and EutQ^100-229^ (Fig. 4C), or for GB1 alone, or for the incubation buffer (Fig. S18). 2D heteronuclear single quantum correlation (HSQC) NMR identified interactions between EutQ^1-99^ and EutC^1-20^, indicated by chemical shift perturbations (CSPs) of selected residues of ^15^N-labelled EutQ^1-99^ in the presence of unlabelled EutC^1-20^ (Fig. 4D). Complete backbone assignment of EutQ^1-99^-GB1 was hampered by severe resonance overlap and/or line-broadening due to the propensity of the H3/H4 region to aggregate; specifically, resonances from the residues 66-77, 82, 85-98 within H3 and H4 regions could not be assigned (Fig. 4E, Fig. S19). The absence of CSPs of residues in H1 and H2 (which were all assigned) confirmed that this region of EutQ is not involved in interactions with EutC^1-20^. Assigned resonances showing CSPs include amino acids 63-65, 78-81, 83, 84 and 99; these are residues found in the linker region and within H3/H4 of EutQ (Fig. 4F). These NMR results confirmed the *in vivo* findings (Fig. 3) and the AlphaFold predictions described above (Fig. S13). We also studied the binding between EutQ and EutE^1-20^, the EP from another cargo protein, EutE, using ITC and NMR (Fig. S20). The EutE binding characteristics, including the binding site and thermodynamics, resembled those observed for EutC^1-20^ binding, albeit with a notable decrease in affinity. EutC^1-20^ and EutE^1-20^ helical peptides show 90% homology with 40% identity (Fig. 4G). Analysis using HeliQuest ^42^ revealed that the conserved amino acids I6, V10, V13 and M17 residues lie on one side of the peptide helices, forming a hydrophobic surface. Surface analyses of the AlphaFold structures of EutQ^1-99^ showed that H3 and H4 together form a hydrophobic crevice (Fig. S13), which provides the binding sites for the EutC and EutE peptides. Hdrophobic contacts were found between the conserved peptide amino acids mentioned above and the I72, L76, F81, L89 and V93 residues of EutQ. The difference in the affinities of EutC and EutE to EutQ could be explained by slight variations in the hydrophilic interactions between the two peptides and EutQ. Together, our results suggest that the EutC^1-20^ and EutE^1-20^ peptides interact with EutQ to mediate cargo encapsulation.

**Fig. 4.**
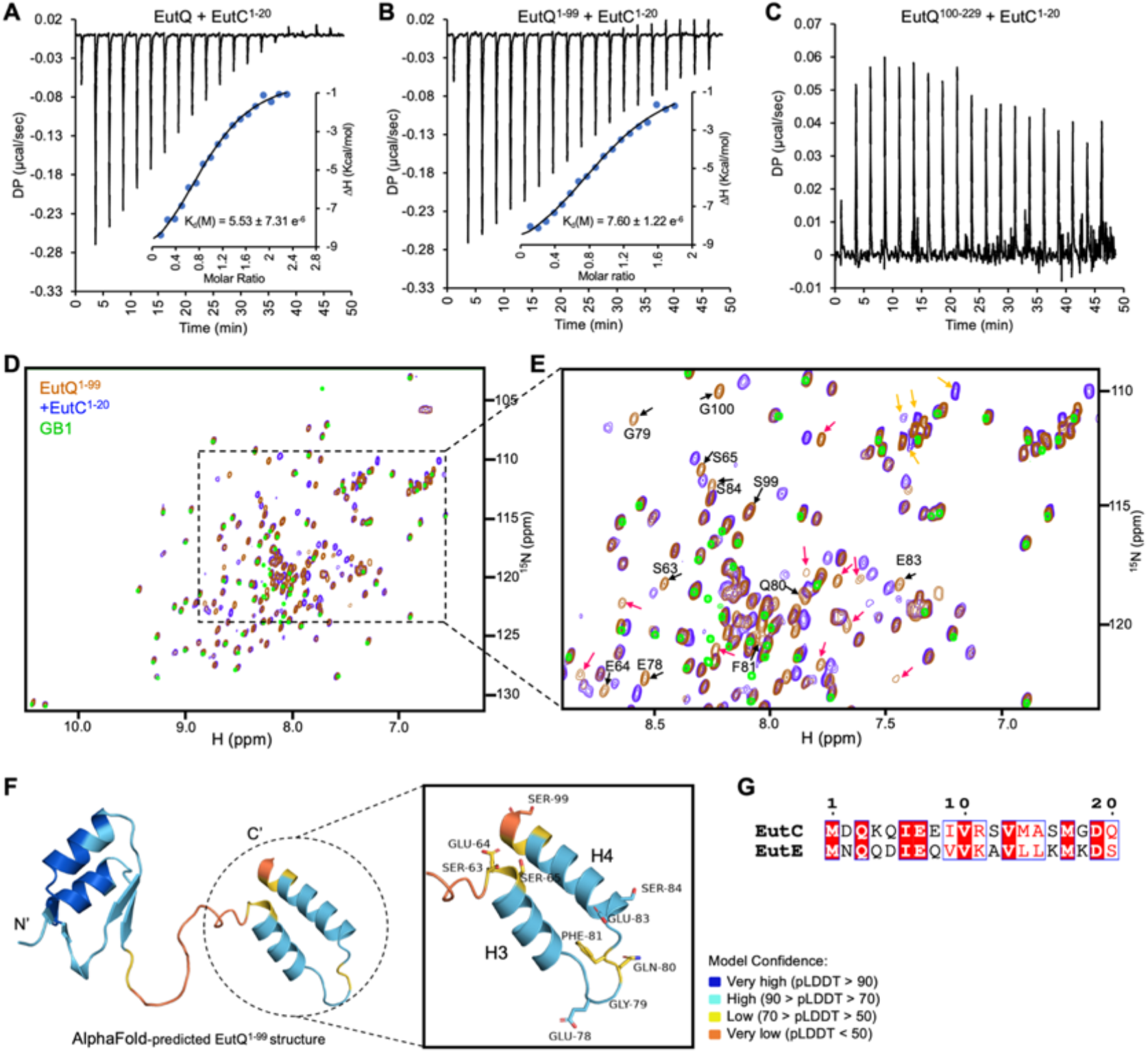
Interactions between EutC^1-20^ and EutQ^1-99^. Fitted isotherm of EutC^1-20^ binding with **A**, EutQ (N=∼2, dimeric EutQ with ligand binding domain contributed by each subunit; ΔG=-7.17 Kcal mol^-1^; ΔH=-10.40 ± 0.51 Kcal mol^-1^; TΔS= -3.24 Kcal mol^-1^), **B**, EutQ^1-99^-GB1 (N=∼1; ΔG=-6.98 Kcal mol^-1^; ΔH=-9.63 ± 0.48 Kcal mol^-1^; TΔS=-2.65 Kcal mol^-1^), and **C**, EutQ^100-229^ (no binding) by ITC. Inset: thermodynamic parameters. Entropic parameters of binding suggest hydrophilic/polar driven interaction between EutQ and EutC^1-20^. **D**, 2D ^15^N-^1^H HSQC spectrum (298K) of ^15^N uniformly labeled EutQ^1-99^-GB1 overlaid with (i) uniformly ^15^N-labeled EutQ^1-99^-GB1 (red) mixed with unlabeled EutC^1-20^ (purple); and (ii) uniformly^15^N-labeled GB1 (green). The chemical shift perturbations (CSPs) observed in EutQ^1-99^-GB1 in the presence of EutC^1-20^ without equivalent shifts on GB1 indicates the specificity of peptide interaction occurring within EutQ^1-99^ alone. **E**, Zoomed-in region of CSPs arising from EutQ^1-99^-GB1 peptide binding. Chemical shift changes occurring in assigned peaks are highlighted with black arrows while those occurring in unassigned peaks are shown in red arrows. Peaks of Gln and Asn side chains that undergo CSPs are designated with yellow arrows. CSP of amino acid residues 63-65, 78-81, 83, 84, and 99 of EutQ^1-99^ in the presence of the peptide were observed while peaks assigned to residues 2-62 remained unchanged. Severe line-broadening of many residues in H3 and H4 helices precluded the specific assignments of many resonances from H3 and H4, limiting the completeness of the data. **F**, AlphaFold-predicted structure of EutQ^1-99^. Resonances that were affected by the presence of EutC^1-20^ are confined to residues from the linker and C-terminal region of EutQ^1-99^. This indicates that interacting region of the EutQ^1-99^ with EutC^1-20^ spans the 63-99 amino acid region. **G**, Sequence alignment of EutC^1-20^ and EutE^1-20^.

Comparative genomic analysis and protein sequence alignment showed that Eut proteins are highly conserved in sequence among Gram-negative bacteria (Figs. S21, S22), implying that the EutQ-mediated assembly mechanisms of Eut BMCs we have discovered in *S*. Typhimurium could serve as a general principle for the assembly of Eut BMCs in a variety of bacterial species. Moreover, because the Eut proteins *S*. Typhimurium and Gram-positive bacteria have a low level of similarity (Figs. S21, S22), we speculate that Gram-positive bacteria employ different Eut BMC assembly mechanisms.

### Eut BMC undergoes a shell-first assembly pathway

To delineate the *de novo* biogenesis of Eut BMCs, we conducted time-lapse live-cell fluorescence imaging to monitor the initial *in vivo* assembly of cargo and shell proteins. We expressed EutM-mCherry/EutB-sfGFP, EutK-mCherry/EutB-sfGFP, or EutB-mCherry/EutM-sfGFP in *S*. Typhimurium cells. After adding EA and B_12_ metabolites to induce expression of native Eut proteins, the first fluorescent spot of Eut shell proteins (indicated by EutM-mCherry, EutK-mCherry, or EutM-sfGFP) emerged from the cytosolic-distribution fluorescence background after ∼30 min of induction. Intriguingly, assembly of shell proteins always appeared prior to assembly of cargo proteins (Fig. 5A, Figs. S23A, S23B). We observed that the first shell assembly tended to occur at the cell pole and along the cell membrane (Fig. 5B). Our results demonstrate that Eut BMCs adopt a unique “Shell first” mode of assembly, a pathway that is distinct from the “Cargo first” assembly of β-carboxysomes ^43,44^ and a combination of “Shell first” and “Cargo first” pathways for Pdu BMCs ^23^ (Figs. S23C, S23D).

**Fig. 5.**
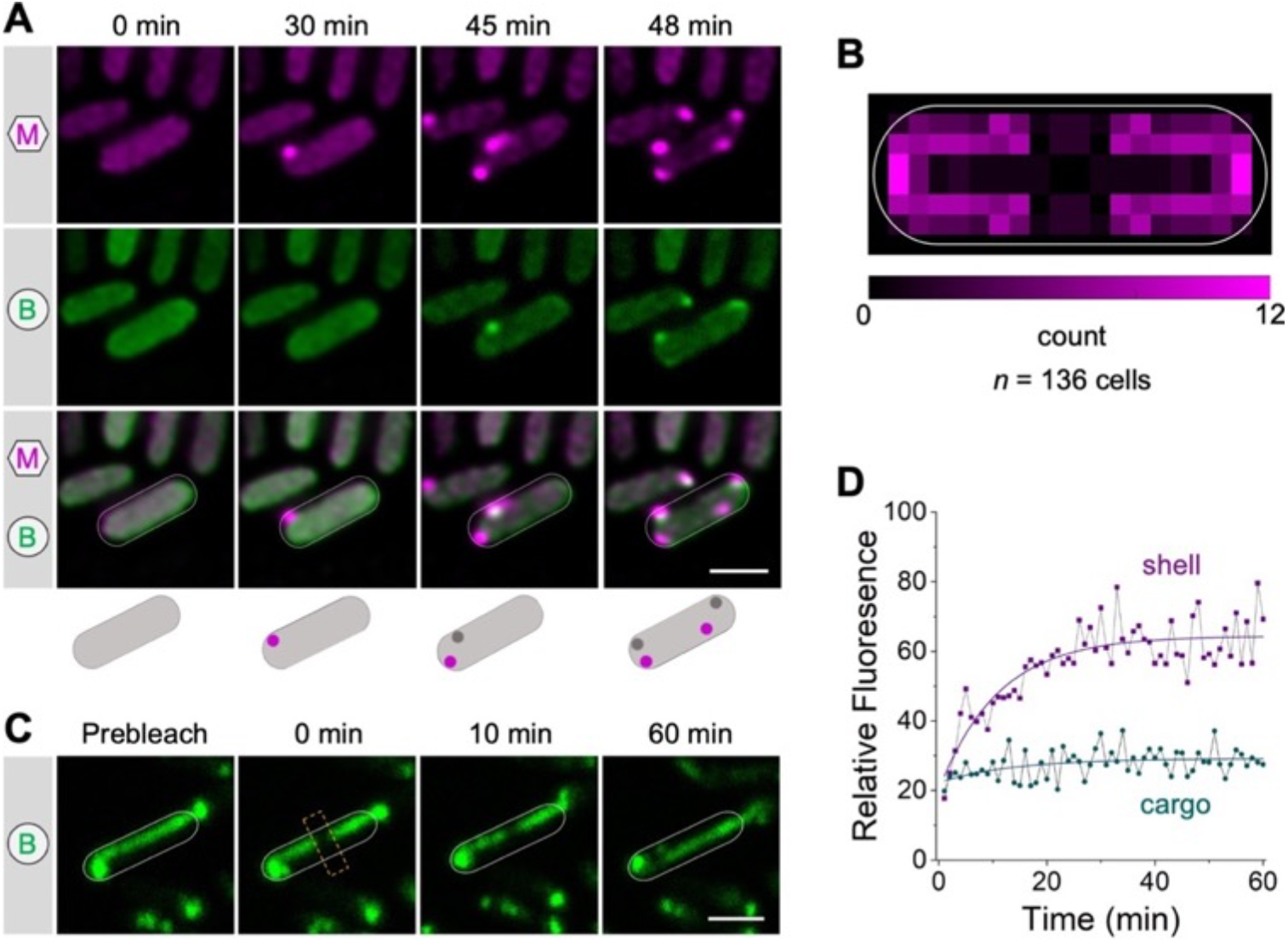
Eut BMC undergoes a “Shell-first” assembly pathway and cargo enzymes form liquid-like condensates within the Eut BMC. **A**, Aggregation of the shell (EutM-mCherry) and cargo (EutB-sfGFP) in the WT strain following induction with EA and B_12_. Scale bar: 1 µm. **B**, Heatmap of the spatial distribution of the first Eut BMC shell within the cell (*n*=136). Scale bar represents the count number of initial shell fluorescence observed at the corresponding location of the cell. **C**, Representative FRAP images on EutB-sfGFP in Δ*eutN* at various time lapses. The yellow rectangular boxes indicated the bleaching area. Scale bar: 1 µm. **D**, Representative time courses of fluorescence recovery of bleached regions of EutB-sfGFP (purple) and EutM-sfGFP (cyan). The y-axis indicates fluorescence values relative to the fluorescence intensity of the selected region prior to bleaching.

### Liquid-like association of cargo enzymes within the Eut BMC

The Eut BMC sequesters a series of cargo protein complexes within the shell. These cargo enzymes can oligomerize via interactions mediated by native EPs, whereas enzymes that lack EPs ‘piggyback’ onto the enzymes containing EPs to achieve encapsulation. To understand the organizational properties of Eut BMCs, we determined the diffusion dynamics of the shell (indicated by EutM) and cargos (indicated by EutB) of Eut BMCs by fluorescence recovery after photobleaching (FRAP). Elongated structures of Eut BMCs formed in the Δ*eutN* strain were used for FRAP experiments and analysis (Fig. 5C, Fig. S24). We found that cargo proteins showed a higher mobile fraction (58.5% ± 4.5%, *n* = 20) than shell proteins (12.8% ± 3.0%, *n* = 20) (Fig. 5D); the half-life of recovery of the cargos was 3.85 ± 2.0 minutes (*n* = 20). We conclude that the cargo proteins form a liquid-like condensate within the Eut BMC, consistent with the findings for α- and β-carboxysomes and for Pdu BMCs of *S*. Typhimurium ^23,45,46^. These findings suggest that oligomerization and condensation of cargo proteins are a widespread phenomenon that occurs within a diverse range of BMCs, which plays a crucial role in BMC assembly and in sustaining their structures and functions.

## DISCUSSION

Eut BMCs are self-assembling proteinaceous organelles that mediate ethanolamine catabolism. Previous attempts using protein crystallization, growth analysis, and synthetic biology had only provided fragmented knowledge about the composition and function of Eut BMCs. The mechanism by which various protein components connect and self-assemble to create a complete and functioning entity remains unclear. Here, we have elucidated the organizational and assembly mechanisms of Eut BMCs in *S*. Typhimurium in unprecedented detail and propose a model for the biogenesis pathway of Eut BMCs (Fig. 6).

**Fig. 6.**
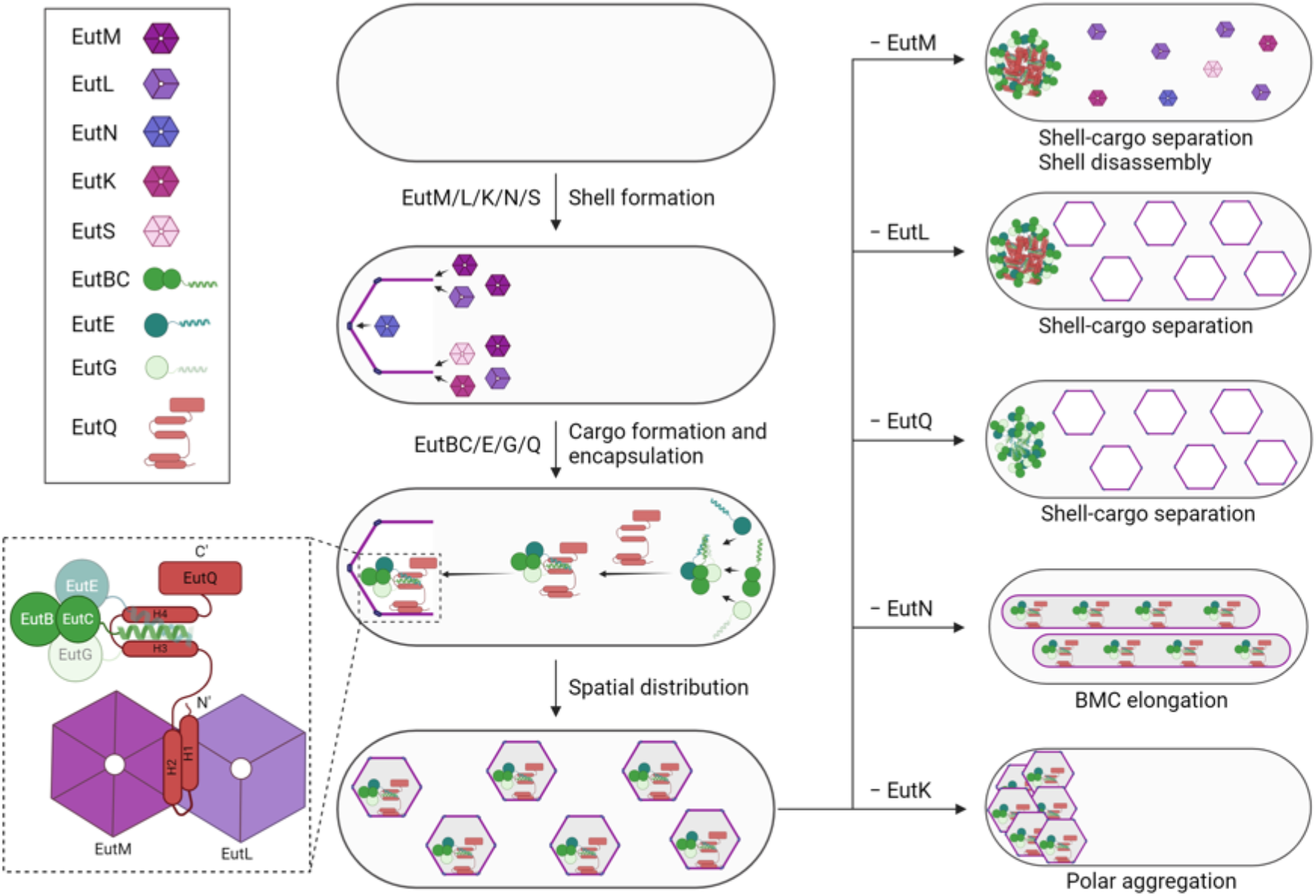
Schematic model of Eut BMC biogenesis. The Eut BMC shell proteins are first recruited by EutM to assemble at a cell pole. Then, cargo proteins self-aggregate at a cell pole and are encapsulated into shell through interactions between EutQ and the N-terminal encapsulation peptides of cargos. EutQ forms a bridge between the cargo and shell proteins: helices H1 and H2 bind the interface between shell proteins, EutL and EutM, whereas cargos are bound to the H3 and H4 helical region, which provides the binding sites for the recruitment of the enzymatic core with shell proteins. The EutN proteins, which likely occupy the vertices, are important the for full encapsulation of individual Eut BMC. EutK is responsible for the subcellular partitioning of Eut BMCs after their biogenesis at the cell poles. EutK is a hexamer due to a single BMC domain at its N-terminus. EutS is a shell component but appears to be not essential for formation of the Eut BMC shell. Created with BioRender.com.

Our data reveal that the Eut BMC undertakes the “Shell-first” assembly pathway, remarkably distinct from the assembly pathways of β-carboxysomes (“Cargo first” pathway) ^43,44^ and Pdu BMCs (concomitant assembly of shell and cargo proteins) ^23^. The assembly of the Eut BMC shell, triggered by EutM, initially occurs at the cell pole. Next, EutQ interacts with the interface of EutM and EutL to provide the binding sites for endogenous EPs of cargo enzymes, enabling cargo encapsulation. In *Salmonella*, the distinct assembly pathways of Eut BMCs and Pdu BMCs might reflect differences in availability of the particular shell and cargo proteins. Analysis of the *Salmonella* transcriptome revealed higher expression levels of genes encoding Eut shell proteins than those encoding Eut cargos, whereas genes encoding *Salmonella* Pdu shell and cargo proteins were expressed at similar levels under anaerobic growth conditions ^47^. Moreover, the assembly of Eut BMC cargos occurs separately from the assembly of its shell, as shell and cargo aggregates were spatially partitioned in the Δ*eutQ* and Δ*eutL* mutants (Fig. 2M). The independent assembly of shell and cargo proteins resembles the observations in Pdu BMCs ^23^. Taken together, our findings indicate that the shell and cargo assembly pathways of different metabolosomes are completely independent, and the temporal sequence of assembly of shell and cargo proteins varies considerably.

We found that the shell proteins, EutM, EutN, EutL, and EutK were essential for the formation and function of Eut BMCs, but EutS was not required. We can draw parallels with the second *Salmonella*-encoded metabolosome, the Pdu BMC. PduU, the EutS homolog within the Pdu BMC, is not required for the assembly and function of Pdu BMCs ^36,48,49^. EutM can self-assemble to form large protein filaments ^50^, which resemble the nano-tube structures formed by the homologous PduA and PduJ proteins ^51-53^. Based on the fact that PduA and PduJ are the two major canonical shell proteins of Pdu BMCs ^48^, we propose that EutM is a major building component of the Eut shell. Our results also show that the pseudohexameric homotrimer EutL ^54^, is essential for the shell-cargo association of Eut BMCs (Fig. 2M). This is consistent with the function of its homolog, PduB, in the assembly of Pdu BMCs ^23^. EutK plays a role in the *in vivo* distribution of Eut BMCs, reminiscent of the role of PduK (with a disordered C-terminal extension) in determining the subcellular locations of Pdu BMCs ^48^. The C-terminal tail of EutK has a positively-charged domain, which could potentially bind to nucleotides ^55^. The molecular mechanism by which EutK governs the partitioning of Eut BMCs and interactions with nucleotides remains to be determined. Overall, our data reveal that the Eut and Pdu shell proteins play similar roles in the assembly of metabolosomes, suggesting that the construction principles of shell proteins we have unraveled may extend to a range of catabolic BMCs in different bacterial hosts.

Our results demonstrate both the structural and catalytic roles of EutQ. Growth experiments suggest that the C-terminus of EutQ has a catalytic function since the absence of EutQ^100-229^ prevented growth when EA was the only available carbon source (Fig. 3D, Fig. S11). The N-terminus of EutQ links the shell and cargos of Eut BMCs, comparable to CcmN, CsoS2, and PduB^1-37^, the linkers proteins that mediate the binding of the shell and cargo enzymes of β-carboxysomes, α-carboxysomes, and Pdu BMCs, respectively ^23,43,46^. Recent studies have revealed that the interactions between CsoS2 (C-terminus) and shell proteins occurred at the interface of shell proteins ^28^, in good agreement with our findings that EutQ could interact with the shell proteins EutL and EutM (Fig. 3I).

## CONCLUSIONS

Our extensive live-cell super-resolution fluorescence imaging, structural, and biochemical studies provide unprecedented insights into the organization and biogenesis of Eut BMCs. We dissected the molecular mechanisms of the stepwise protein assembly and the hierarchical cooperation that governs the formation of structurally defined and functional Eut BMCs. This breakthrough advances our knowledge of catabolic BMC assembly and provides a molecular roadmap for reprogramming and modulating BMCs to improve the efficiency of metabolic reactions. The future identification of specific binding sites for BMC shell and cargo protein interactions could pave the way for the development of therapeutic interventions for a variety of infectious diseases that target the mammalian gastrointestinal tract.

## METHODS

### Bacterial strains and growth conditions

The bacterial strains used in this study derived from *S. enterica* subsp. *enterica* serovar Typhimurium LT2 (*S*. Typhimurium LT2) ^56^. The rich medium was LB Broth (Lennox) medium (10 g·L^-1^ tryptone, 5 g·L^-1^ yeast extract, 5 g·L^-1^ sodium chloride), and the minimal medium was no-carbon-E (NCE) medium supplemented with 1 mM MgSO_4_, 0.5% succinate, and 30 mM ethanolamine and 200 nM vitamin B_12_ (if applicable). LB plate was prepared with 15 g·L^-1^ agar, and minimal medium plate was prepared with 2% (w/v) low-melting point agarose. Antibiotics were added to media as required at the following final concentrations: ampicillin at 100 μg·mL^-1^, kanamycin at 50 μg·mL^-1^, and gentamicin at 20 μg·mL^-1^ in ddH_2_O, chloramphenicol at 25 μg·mL^-1^ in ethanol, and tetracycline at 25 μg·mL^-1^ in methanol.

Intracellular visualization of fluorescently-tagged Eut proteins was conducted following the growth of the plasmid-carrying strains in minimal medium under aerobic conditions. An overnight LB culture was inoculated 1:100 in 100 μL of minimal medium in a 2-mL Eppendorf tube in the absence of EA and B_12_ shaken horizontally at 220 rpm overnight. Unless otherwise specified, 1 μL of this culture was sub-inoculated to 100 μL of minimal medium in a 2-mL Eppendorf tube, both in the absence and in the presence of EA and B_12_, shaken aerobically at 37°C for 5 hours. For birth event detection, 5 μL of the sub-inoculated culture (in minimal medium without EA and B_12_) was dropped onto minimal medium plate in the presence of EA and B_12_ and incubated aerobically at 37°C for time-lapse imaging.

### Construction of chromosomal mutations

All mutants in this study were generated by scar-less genome editing technique developed previously using the pEMG and pSW-2 plasmids ^57^ (Fig. S4). A pair of DNA fragments (∼800 bp) flanking the chromosome regions of interest were firstly PCR amplified and integrated into the linear pEMG suicide plasmid (digested by EcoRI and BamHI) by Gibson assembly (NEBuilder HiFi DNA Assembly kit) ^58^. The pEMG-derivative plasmids were mobilized from *E. coli* S17-1 λ *pir* to *S*. Typhimurium by conjugation. *S*. Typhimurium transconjugants that have integrated pEMG-derivative plasmids were selected on M9 minimal medium plates supplemented with kanamycin and 0.2% of glucose. pSW-2, extracted from *S*. Typhimurium LT2, was subsequently transformed into the transconjugants by electroporation. Then, transformants were selected on LB plates with gentamicin and 1 mM of *m*-toluate added. Colonies were screened for kanamycin resistance and sensitive clones were verified by PCR. pSW-2 was finally cured from the resulting strains by two passages in LB without adding gentamicin. Please note, for the deletion of N-terminus of EutQ, the first nine amino acids were kept due to an overlap of coding genes between *eutP* and *eutQ*. Strains and plasmids used in this study are listed in Table S1. A complete list of primers can be found in Table S2. All mutants were verified by PCR and DNA sequencing of PCR-amplified genomic DNA sequencing (Fig. S5).

### Construction of vectors

pBAD/*Myc*-His was used as a backbone for generation of fluorescence-tagging vectors. pBAD/*Myc*-His was first digested by NcoI and HindIII. Then PCR-amplified DNA fragments of individual *eut* genes, mCherry, and sfGFP were integrated into the linear vector by Gibson assembly ^58^. pXG10-SF containing a constitutive P_LtetO-1_ promoter was employed as a backbone for complementation experiments ^59^. The PCR-amplified coding sequences of EutK/L/Q were inserted into linear pXG10-SF (obtained by PCR cloning) via Gibson assembly. Plasmids and primers used in this study were listed in Table S1 and S2, respectively. All constructed vectors were verified by PCR and plasmid sequencing.

### Bacterial growth assays

The *S*. Typhimurium strains of interest were firstly inoculated from isolated colonies from LB plates into 5 mL of LB in 30 mL universal glass vials, and then incubated at 37°C overnight with shaking at 220 rpm. The overnight cells were washed three times and then resuspended in M9 medium (supplemented with 2 mM MgSO_4_, 100 μM CaCl_2_, 30 mM ethanolamine, and 200 nM B_12_) to an OD_600_ of 0.01. 250 μL of the culture was growing aerobically at 37°C in a 96-well microplate reader with shaking at 225 rpm, using the System Duetz technology platform (Growth Profiler 960, EnzyScreen). The readings of OD_600_ were taken every 30 min. At least three biological replicates of each growth curve were collected. Lag time was calculated by time to reach OD_600_=0.2. Non-linear regression was used for calculation by the Gompertz Growth model.

### Transmission electron microscopy

The structures in the *S*. Typhimurium WT and mutant strains were visualized using thin-section electron microscopy. An overnight LB culture was inoculated 1:100 in 5 mL of minimal medium in a 30 mL universal glass vial in the absence of EA and B_12_ shaken at 220 rpm overnight at 37°C. 500 μL of this culture was sub-inoculated to 50 mL of minimal medium in a 250 mL flask in the presence of EA and B_12_, shaken aerobically at 37°C for 5 hours. Bacterial samples were pelleted at 6,000 g for 10 minutes and fixed with 0.05 M sodium cacodylate buffer (pH 7.2) containing 2.5% glutaraldehyde using 100 W for 1 minute twice (P1). Samples were embedded in 4% agarose, and then stained with 2% osmium tetroxide and 3% Potassium Ferrocyanide using 100 W for 20 seconds for three times (P2). The reduced osmium stain was then set using a 1% Thiocarbohydrazide solution for 10 minutes. The second osmium stain was applied using P2 with 2% osmium tetroxide. The sample was made electron dense with 2% Uranyl Acetate incubated at 4°C overnight. Dehydration was run with a series of increasing alcohol concentrations (30% to 100%) before cells embedding in medium resin. Thin sections of 70 nm were cut with a diamond knife. Images were recorded by a Gatan Rio 16 camera, DigitalMicrograph software, and an FEI 120Kv Tecnai G2 Spirit BioTWIN transmission electron microscope.

### Super-resolution fluorescence microscopy

The bacterial cells were imaged using a ZEISS Elyra 7 with Lattice SIM2 super-resolution microscope. Cells were pelleted and washed twice with PBS buffer, followed by fixing with 4% of paraformaldehyde solution (prepared with PBS buffer) and incubated at room temperature for 15 minutes before imaging. Images were captured under Lattice SIM using SBS LP 560 dual camera beam splitter and a Plan-Apochromat 63×/1.4 oil DIC M27 objective. Two tracks (one for sfGFP imaging excited at 488 nm and the other one for mCherry imaging excited at 561 nm) detected by two cameras were used to avoid fluorescence cross-talking. Two cameras were always aligned at calibration mode before imaging. Images were captured as 1280 x 1280 pixels at 16 bits. Super-resolution images were obtained by processing under SIM^2^ by Zen software. To monitor the birth event and the progression of the shell and the cargo of Eut BMCs, 5 μL of bacterial culture in minimal medium in the absence of EA and B_12_ was dropped onto minimal medium plate in the presence of EA and B_12_ and left to dry at 37°C. The agraose plate with cell patches was cut out and attached to a 35mm glass-bottom dish and covered by a 0.17 mm glass coverslip. The images were taken at 0 min when cells just started to grow on minimal medium plate in the presence of EA and B_12_, and every three minutes for tacking the assembly of Eut BMCs. Represented images were shown at different time points when important events occurred. The temperature was controlled at 37°C during imaging. All images were captured from at least three biological replicates.

Fluorescence recovery after photobleaching experiments were performed on a Zeiss LSM780 confocal microscopy as described previously ^23^. 5 μL of bacterial cells that grew in minimal medium in the presence of EA and B_12_ for 5 hours were dropped onto minimal medium plate and allowed to dry before sandwiching with a 35mm glass-bottom dish and a 0.17 mm glass coverslip. 100% laser power was applied to bleach a line across the center of the cell. Images were captured every 1 min for 60 min. Fluorescence profiles along the long axis of the cell were obtained by ImageJ software and normalized to the same total fluorescence. The mobile proportion (M) was given by:

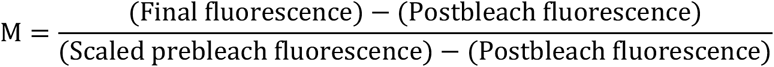

Experimental fluorescence recovery curves were obtained by plotting the fluorescence profile values at the center of the bleached area against time. Then fluorescence recovery curves were fitted to a single exponential function, given by:

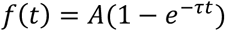

### Protein expression and isolation

The encapsulation peptides EutC^1-20^ and EutE^1-20^ were synthesized by GenicBio. Synthetic gene for EutQ (residues 1-229, Uniprot code: Q9ZFV5) was cloned into pET14b modified with a TEV protease site located prior to the protein gene sequence. EutQ N-terminal domain fused with a soluble fusion partner, the immunoglobulin binding domain of streptococcal protein G (GB1) at its C terminus (EutQ^1-99^-GB1), EutQ C-terminal domain (EutQ^100-229^) and GB1 were cloned into pETM11 expression vector. Hexahistidine Ni^2+^ affinity tag (His-tag) was positioned between EutQ^1-99^ and GB1. Vector backbone and primers for Gibson assembly cloning are listed in Tables S1 and S2.

Heterologous expression of recombinant EutQ, EutQ^1-99^-GB1, EutQ^100-229^, and GB1 were achieved by transforming BL21 (DE3) star cells with the relevant expression plasmid. Cells were plated and incubated overnight on LB agar containing either Ampicillin (pET14b) or Kanamycin (pETM11) at 37°C in a static incubator. Starting *E. coli* culture were prepared by inoculating a 1 mL LB broth containing the relevant antibiotics was inoculated with 1 colony picked from the prepared plates and incubating at 210 rpm for 5 hours, at 37°C. For protein expression in minimal media, another starting *E. coli* culture were prepared by inoculating 20 mL M9 media with 100 μL primary culture in the presence of the relevant antibiotics and incubated overnight at 210 rpm, 37°C. M9 protein expression media for preparing the ^15^N and ^13^C isotopically-labelled proteins were prepared with either isotopically labeled ^15^NH_4_Cl for 2D NMR or with both ^15^NH_4_Cl and ^13^C-glucose for 3D NMR studies. Expression culture was then made by adding 20 mL starting culture into 1 L M9 media containing the relevant antibiotics and incubated at 37°C, 210 rpm until the OD_600_ of 0.7 was reached.

Protein expression was induced with 0.8 mM IPTG. The cell culture was incubated overnight at 18°C, 180 rpm, and cells were harvested by centrifugation at 5,000 rpm for 30 mins at 4°C. Cell pellet of 2 L culture were resuspended to 5% w/v in buffer A (50 mM Na_2_HPO_4_, 300 mM NaCl, 1 mM DTT, 10% glycerol pH 7.2) supplemented with 15 μg·mL^-1^ DNase1, stood in ice for 20 mins and lysed by one pass through the cell disruptor (Constant Systems, UK) at 19 KPSI at 4°C. Lysate was centrifuged at 18000 rpm, 4°C and the supernatant collected and passed through a pre-equilibrated 5 mL FF His-trap column (Cytiva, USA). The column was washed with 5 column volumes (CV) of 5% buffer B (buffer A + 500 mM imidazole) and eluted by a gradient increase of buffer B. Protein samples were further purified by size exclusion chromatography using S200 Superdex 16/600 GE healthcare column in buffer containing 50 mM Na_2_HPO_4_,150 mM NaCl, 10% glycerol, pH 7.2. These were aliquoted and stored in -80°C. Experimental protein samples were buffer exchanged into 20 mM Na_2_HPO_4_, 20 mM NaCl, pH 6.5.

### NMR Spectroscopy

Uniformly ^15^N labeled recombinant EutQ^1-99^-GB1 used was diluted to a final concentration of 100 µM of 600 µL NMR buffer (20 mM Na_2_HPO_4_, pH6.5; 20 mM NaCl) supplemented with 10% v/v D_2_O. Binding studies was done by mixing ^15^N-labelled EutQ^1-99^-GB1 or EutQ with unlabeled peptides (either EutC^1-20^ or EutE^1-20^) in a 1:5 molar ratio of EutQ:peptide. All ^15^N-H HSQC NMR experiments were acquired at 298K on Bruker Avance III 700MHz spectrometer equipped with TCI cryoprobe [^1^H, ^15^N, ^13^C] with frequency locked on ^2^H_2_O. Water suppression was achieved by excitation sculpting. Protein backbone resonance assignment of uniformly ^15^N, ^13^C-labeled 1mM EutQ^1-99^-GB1 was achieved using 25% non-uniform sampling of three pairs of 3D Triple resonance experiments – CBCA(CO)NH/CBCANH; HBHA(CO)NH/HBHANH; and HNCO/HN(CA)CO. Data analysis was performed with CcpNMR Analysis version 3.1.1.

### Isothermal titration calorimetry (ITC)

Malvern Microcal Peaq Automated fitted with its proprietary software (MicroCal PEAQ_ITC Automated Control Software) was used for the experiment. The instrument was set for 19 injections at 25°C while reference power, initial delay and stir speed were set for 6ucal/s, 60s and 750 rpm respectively. Cell and syringe contained, respectively, 20 μM protein (EutQ/EutQ^1-99^-GB1/EutQ^100-229^/GB1) and 200-400 μM encapsulation peptides. Peptide to buffer titration was used as the internal control. Both protein and ligand were diluted in buffers containing 20 mM Na_2_HPO_4_; 20mM NaCl pH6.5. Start injection volume was 0.4 μL while subsequent injections were 2 μL. MicroCal PEAQ-ITC Analysis Software was used for data evaluation.

### Bioinformatics analysis

According to previous studies, bacteria from 10 genera were reported to contain the *eut* operon: *Citrobacter, Clostridium, Enterococcus, Escherichia, Klebsiella, Listeria, Proteus, Salmonella, Shigella, Yersinia* ^60,61^. To screen the bacterial species that produce the Eut BMCs, the representative genomes of bacterial species from the 10 genera were downloaded from RefSeq database (ftp://ftp.ncbi.nlm.nih.gov/genomes/). A protein BLAST database was made from the translated CDS of the representative genomes. The protein sequences of the *eut* operon from *S*. Typhimurium LT2 (RefSeq ID: ASM694v2) were queried against the BLAST database using BLASTp v2.5.0+ ^62^. According to the presence of the essential proteins including EutB, EutL, and EutM, 30 representative isolates were found to contain the Eut BMCs (Supplemental File 1).

To infer the phylogenetic relationship of the 30 bacterial isolates, the core genes of the isolates were extracted using BUSCO v5.2.5 ^63^ with “bacteria_odb10” database. From the single-copy orthologs detected by BUSCO, 113 proteins were existing in all the genomes. The orthologs were extracted and then aligned using Mafft v7.475 ^64^. The alignments were then merged using Seqkit v0.15.0 ^65^ and trimmed with trimAl ^66^. Fasttree v2.1.10 ^67^ was used to infer the phylogenetic tree using the JTT + Gamma model. The tree was mid-rooted. The identity of Eut proteins summarized from the BLASTp result was visualized with the tree in iTOL ^68^. The sequences of EutQ, EutC, and EutE proteins were aligned by ESPript 3 ^69^.

## AUTHOR CONTRIBUTIONS

M.Y., O.A., P.C., X.Z., Y.L., G.F.D., Y.C. and N.S. performed research; M.Y., O.A. and L.-N.L. analyzed data; M.Y., O.A., J.C.D.H, L.-Y.L. and L.-N.L. designed research; M.Y., O.A., J.C.D.H, L.-Y.L. and L.-N.L. wrote the article with contributions from other authors.

