## Supplementary Information for "Molecular basis of the biogenesis of a protein organelle for ethanolamine utilization"

### Supplemental Information

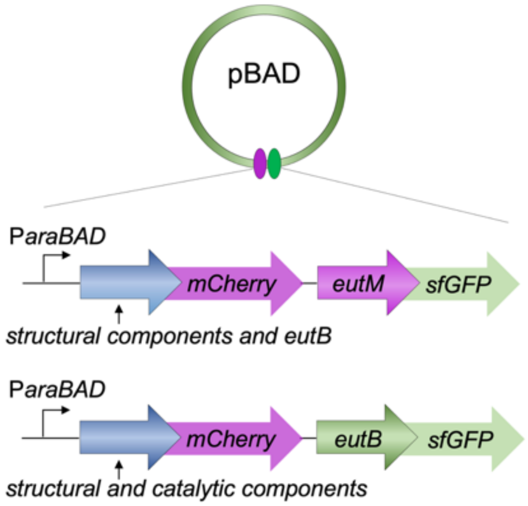


**Figure S1. Constructed vectors to visualize the locations of Eut proteins.** Eut proteins are visible by dual-labeling with fluorescence proteins (mCherry and sfGFP). Two series of vectors were constructed. First, EutM (a shell protein) was labeled with sfGFP, and other structural components or EutB (an enzymatic component) were labeled with mCherry. Second, EutB was labeled with sfGFP, and the structural components or other catalytic components were labeled with mCherry.

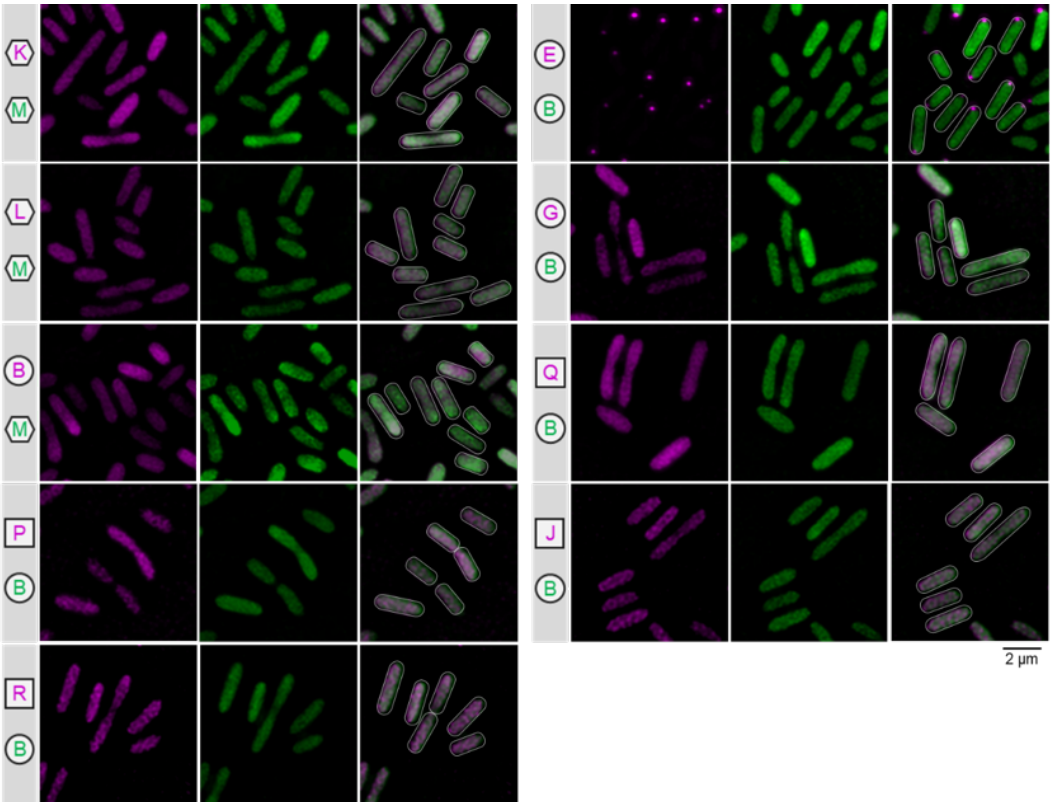


**Figure S2. Eut BMCs are not formed in the absence of EA and $B_{12}$.** *S.* Typhimurium carrying pBAD to express Eut proteins tagged with mCherry and sfGFP, was growing in minimal medium in the absence of EA and $B_{12}$. EutE aggregated at the cell pole, likely due to its encapsulation peptides at the N-terminus (Quin et al., 2016).

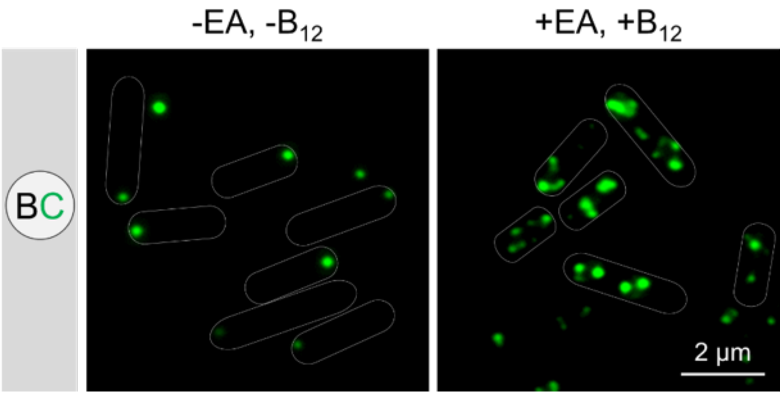


**Figure S3. The distribution of EutBC in the absence and presence of EA and $B_{12}$.** *S.* Typhimurium carrying pBAD to express EutB and EutC-sfGFP, was growing in minimal medium in the absence and presence of EA and $B_{12}$. EutBC aggregated at the cell pole in the absence of EA and $B_{12}$, likely due to its encapsulation peptides at the N-terminus (Choudhary et al., 2012).

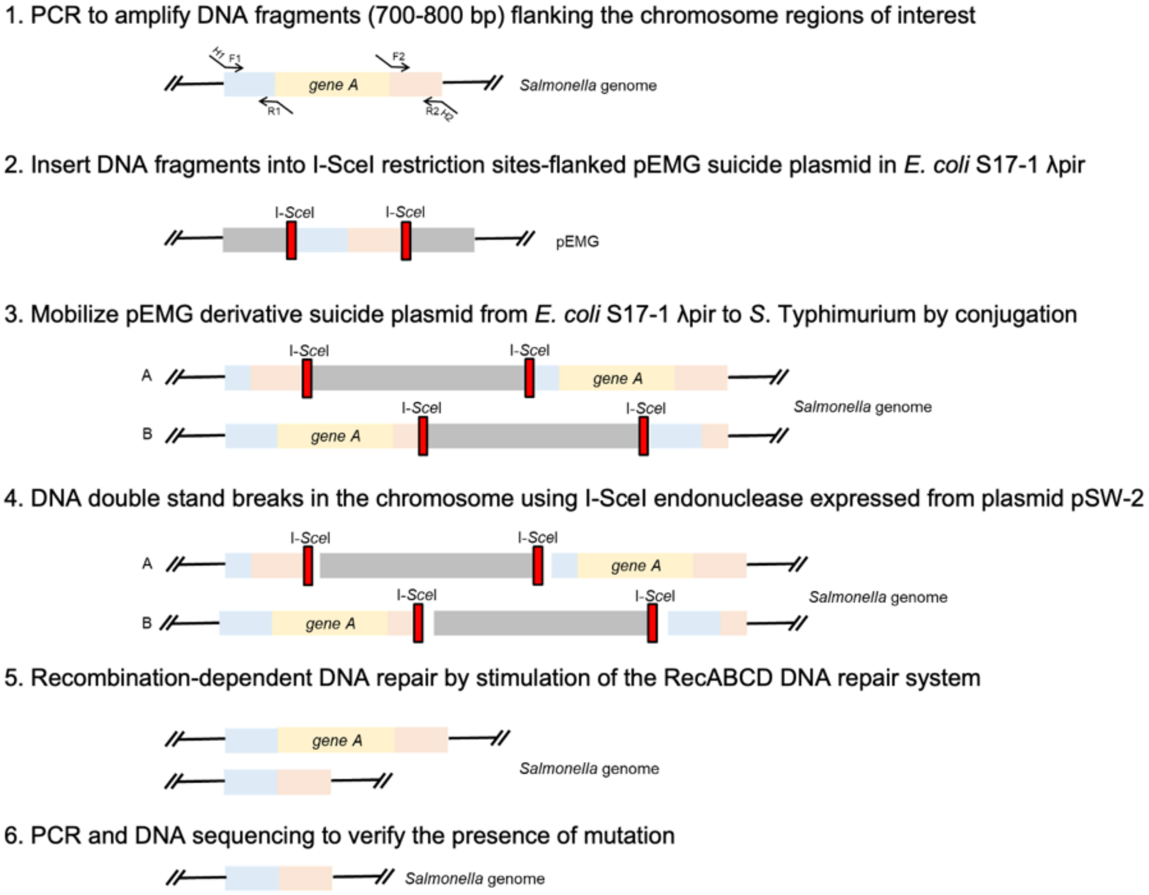


**Figure S4. Workflow for construction of gene deletion mutants for *eut* genes in *S.* Typhimurium**. H1 and H2 refer to the homology extensions or regions of pEMG. F1, R1, F2 and R2 refer to priming sites.

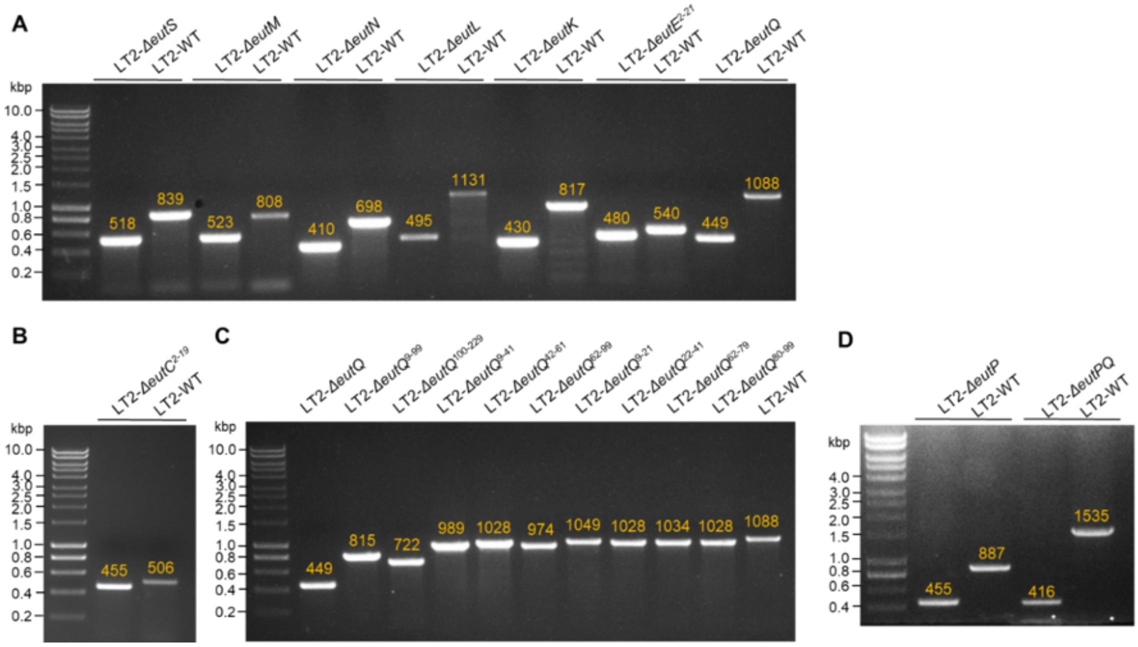


**Figure S5. PCR-based verification of genome editing mutants. A**, *eut* shell gene deletion mutants, Δ*euE*$^{2\text{-}21}$ and Δ*eutQ*. **B**, Δ*eutC*$^{2\text{-}19}$. **C**, *eutQ* related gene deletion mutants. **D**, Δ*eutP* and Δ*eutPQ*. The sizes of the PCR products are indicated (bp, yellow).

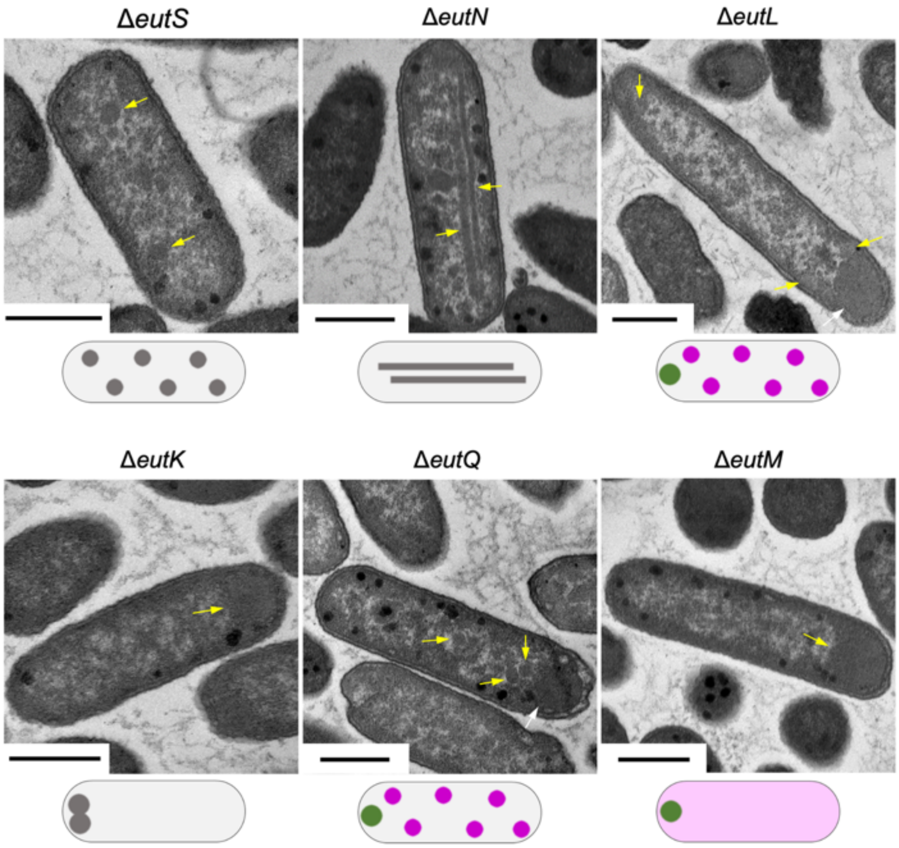


**Figure S6. Thin-section EM verified the observations from the fluorescence imaging.** The strains used for thin-section EM were not transformed with any fluorescent-labeled vectors. Schematic models of the *in vivo* localization of shell and cargo assemblies observed from fluorescence imaging (Figs. 2**A-F**) were shown at the bottom for comparison. Scale bar: 500 nm.

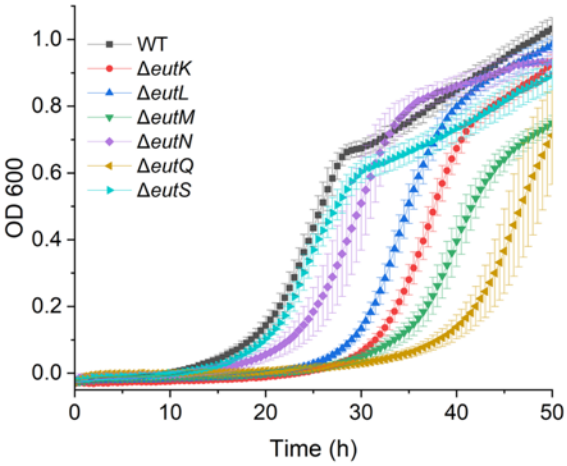


**Figure S7. Growth curves of the *S*. Typhimurium WT and the *eut* gene deletion mutants.** Cells were grown under aerobic conditions in M9 medium, which contains 30 mM EA, 200 nM vitamin $B_{12}$, 2 mM $MgSO_4$, and 100 µM $CaCl_2$.

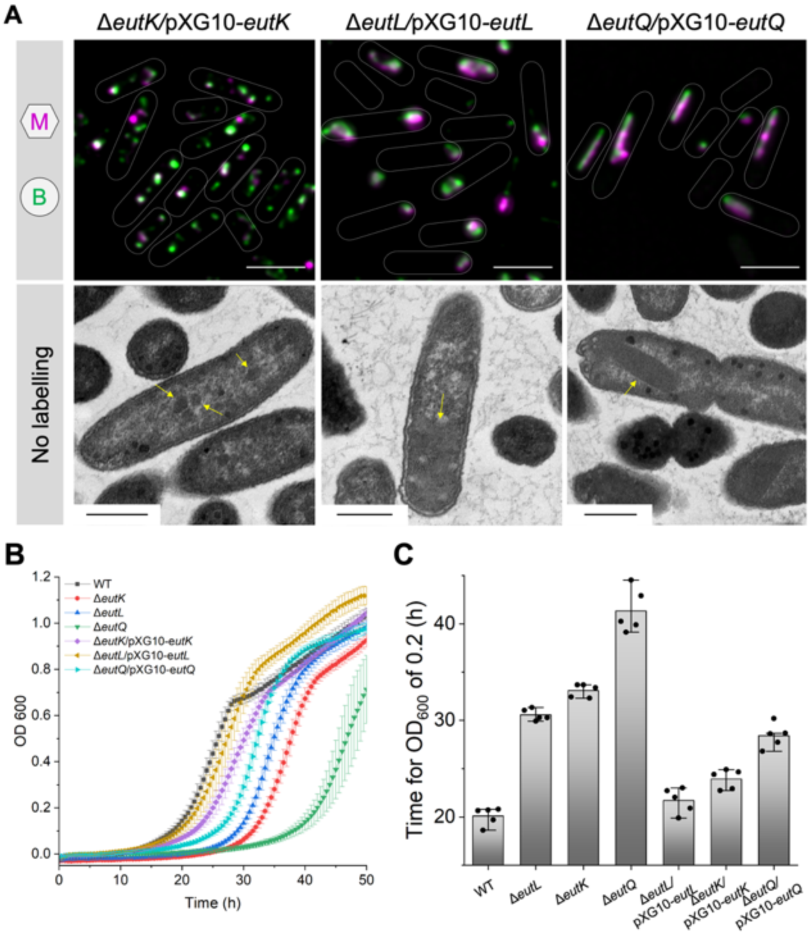


**Figure S8. Successful complementation of EutK, and partial complementation of EutL and EutQ. A**, Fluorescence imaging and thin-section EM on gene deletion mutants expressing deleted proteins from pXG10 plasmid growing in minimal medium in the presence of EA and $B_{12}$. Scale bar in fluorescence images: 2 µm. The Eut BMC structures are indicated with yellow arrows. Scale bar in EM images: 500 nm. **B**, Growth assay curves on gene deletion mutants expressing deleted proteins from pXG10 plasmid grown on EA in M9 medium compared with WT and related mutants. **C**, Time for LT2 WT and mutants to grow to $OD_{600}$=0.2 on EA and $B_{12}$ in M9 medium ($n$ = 5). The center for error bars represents the mean. The whiskers extend to the smallest and largest data points that are within 1.5 times the interquartile range of the upper and lower quartiles. $n$ number of biologically independent experiments.

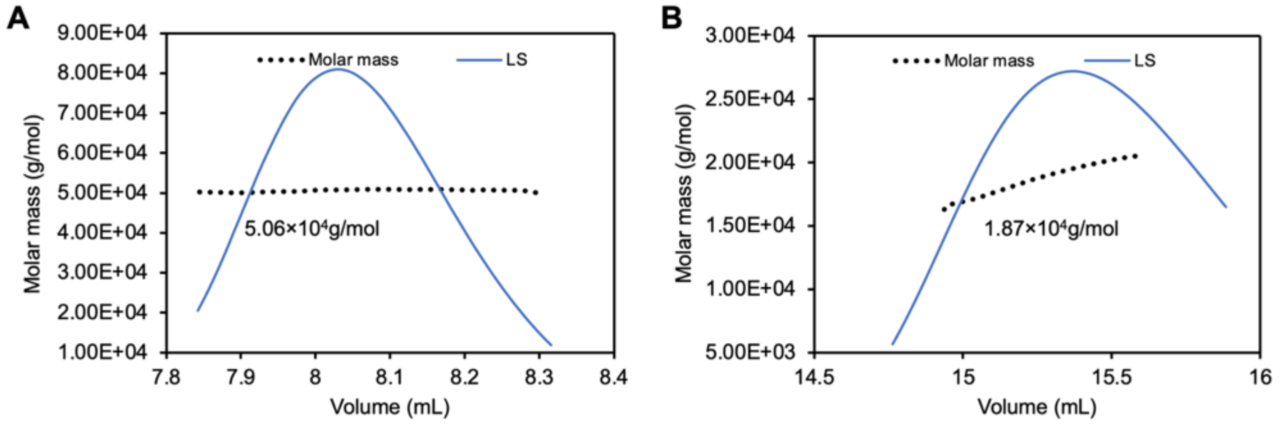


**Figure S9**. **Molecular mass analysis by SEC-MALS with the fitting trace presented. A**, EutQ analysis by SEC-MALS revealed a ~50 KDa protein dimer (the molecular weight of a monomer is ~25 KDa). **B**, SEC-MALS analysis of EutQ$^{1\text{-}99}$ fused with GB1 revealed a ~19 KDa protein (expected molecular weight: 19.64 KDa), indicating that the dimerization of EutQ was mediated by the contacts between two EutQ C-termini.

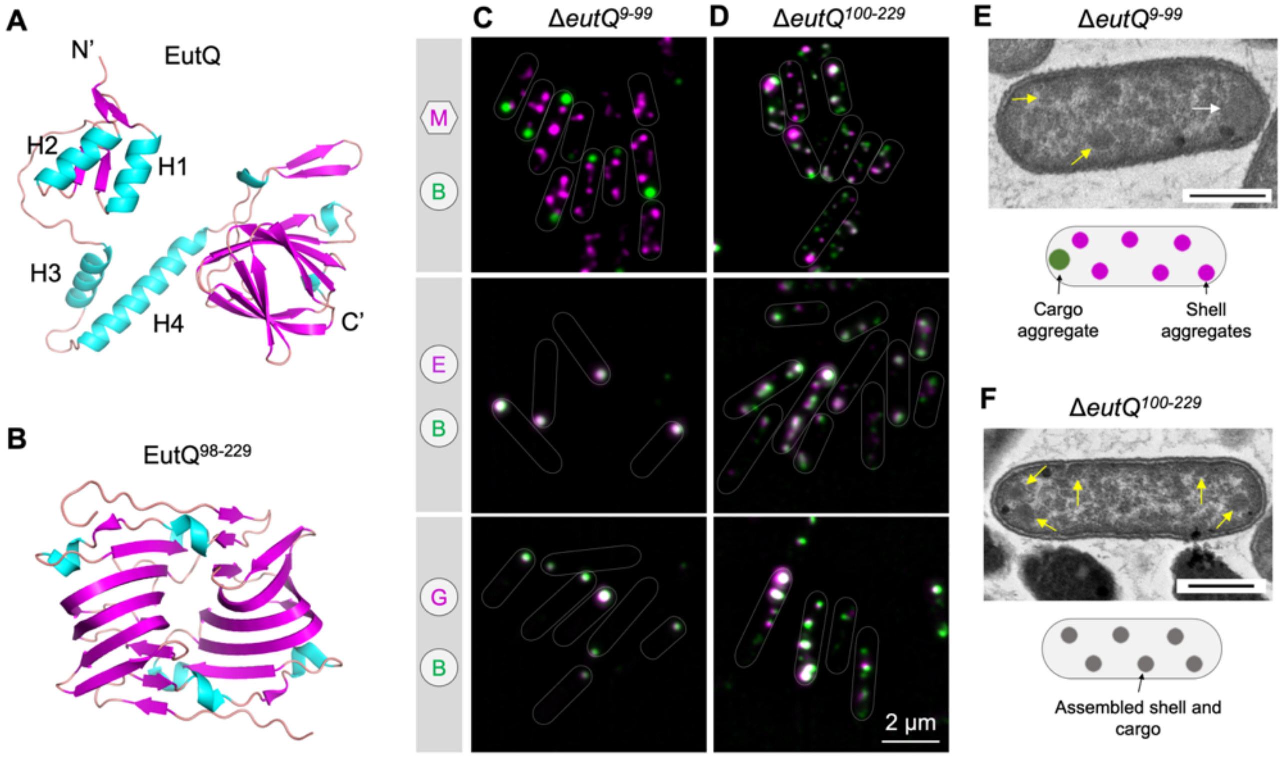


**Figure S10. The N-terminus of EutQ binds the enzymatic core to the shell of EutBMC. A**, The structure of EutQ predicted by AlphaFold. **B**, The crystal structure of the C-terminus of EutQ (PDB ID: 2PYT). **C** and **D**, The location of shell protein EutM and different catalytic components were visualized Δ*eutQ*$^{9\text{-}99}$ and Δ*eutQ*$^{100\text{-}229}$ growing in the presence of EA and $B_{12}$. **E** and **F**, Thin section EM on Δ*eutQ*$^{9\text{-}99}$ and Δ*eutQ*$^{100\text{-}229}$ growing in the presence of EA and $B_{12}$. Schematic models of the *in vivo* localization of shell and cargo assemblies observed from fluorescence imaging (Figs. 3B, 3C, Figs S10C, S10D) were shown at the bottom for comparison. White arrow indicates polar aggregate. Scale bar: 500 nm.

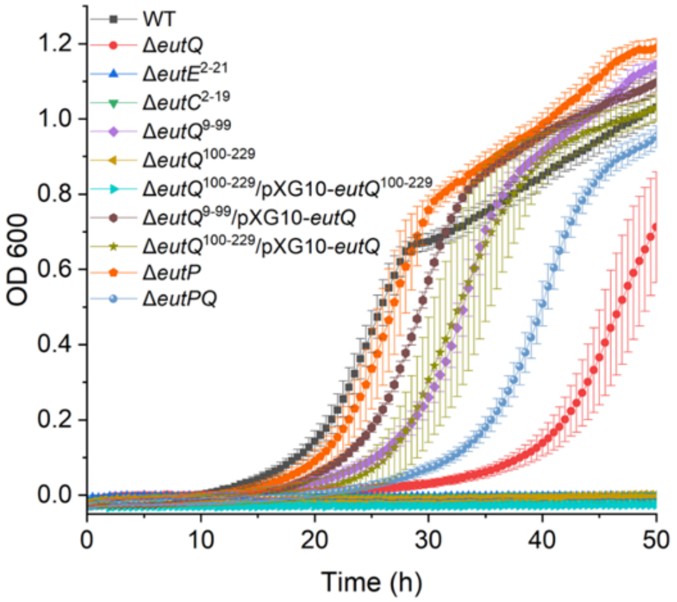


**Figure S11. Growth curves of the *S*. Typhimurium WT, the encapsulation peptides deletion mutants, and *eutQ*-related mutants.** The medium is M9 medium, containing 30 mM EA, 200 nM vitamin $B_{12}$, 2 mM $MgSO_4$, and 100 μM $CaCl_2$. Growth curves were measured on a Growth Profiler 960 (EnzyScreen) under aerobic conditions.

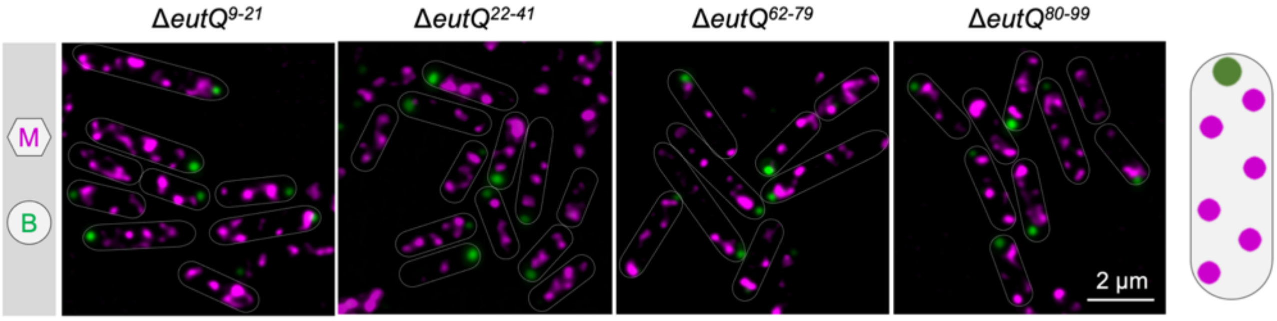


**Figure S12. Individual alpha-helix at EutQ N-terminus are essential for the assembly of shell and cargo of Eut BMCs.** EutM-mCherry (shell) and EutB-sfGFP (cargo) were visualized in individual alpha-helix deletion mutants at EutQ N-terminus, following growth in the presence of EA and $B_{12}$.

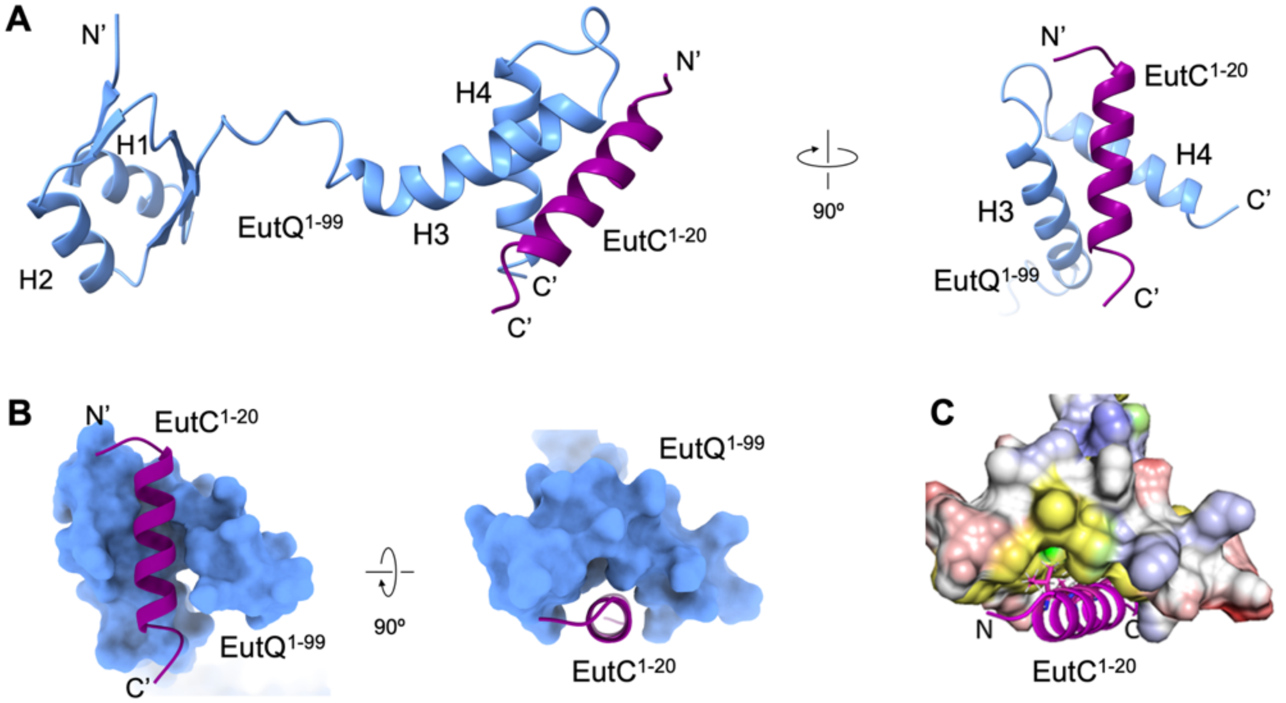


**Figure S13. Interactions between EutQ N-terminus and EutC$^{1-20}$ predicted by AlphaFold. A**, EutC$^{1-20}$ binds to the H3 and H4 α-helices of EutQ$^{1-99}$. **B,** Surface presentation of the binding pocket formed by the H3 and H4 α-helices of EutQ$^{1-99}$ to EutC$^{1-20}$. **C**, Electrostatic potential of the binding pocket formed by the EutQ$^{1-99}$ H3 and H4 α-helices. Hydrophobic residues I72, L76, F81, L89 and V93 (yellow surface) of EutQ form the main contacts with residues I6, V10, V13 and M17 (stick representation) of EutC. The electrostatic surface was calculated using eF-Surf (https://pdbj.org/eF-surf/top.do).

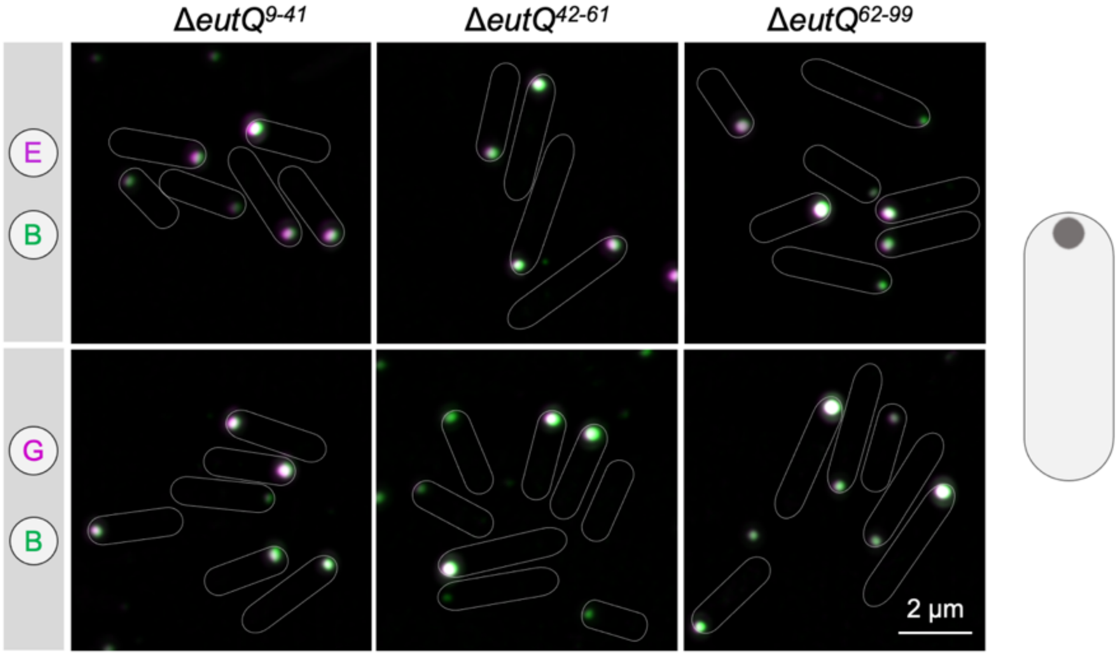


**Figure S14. The location of the cargo proteins of Eut BMCs in different gene deletion mutants of *eutQ*.** EutE-mCherry/EutB-sfGFP and EutG-mCherry/EutB-sfGFP were visualized in different mutants following growth in minimal medium in the presence of EA and $B_{12}$.

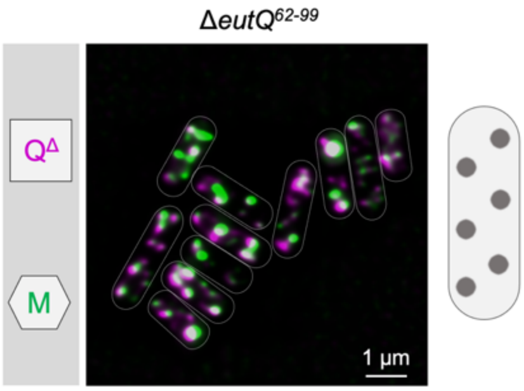


**Figure S15. EutQ$^{\Delta 62\text{-}99}$ assembled with the shell in Δ*eutQ*$^{62\text{-}99}$.** EutQ$^{\Delta 62\text{-}99}$-mCherry and EutM-sfGFP (shell) were visualized in Δ*eutQ*$^{62\text{-}99}$ in the presence of EA and $B_{12}$.

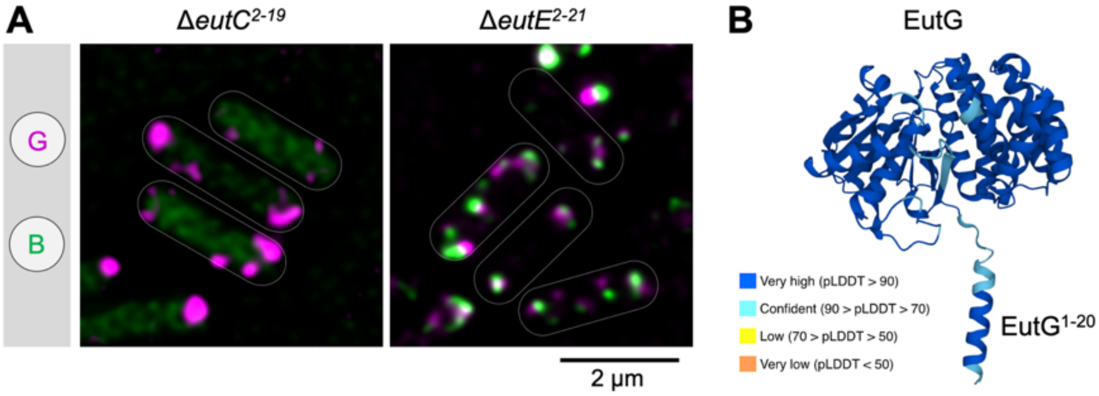


**Figure S16. EutG may possess its own encapsulation peptide for recruitment into the Eut BMC. A,** The location of EutG in EP-deletion mutants. EutG-mCherry and EutB-sfGFP were visualized in Δ*eutC*$^{2\text{-}19}$ and Δ*eutE*$^{2\text{-}21}$ following growth in minimal medium in the presence of EA and $B_{12}$. **B**, AlphaFold predicted structures of EutG and its potential EP, EutG$^{1\text{-}20}$ (Uniprot ID: P41795).

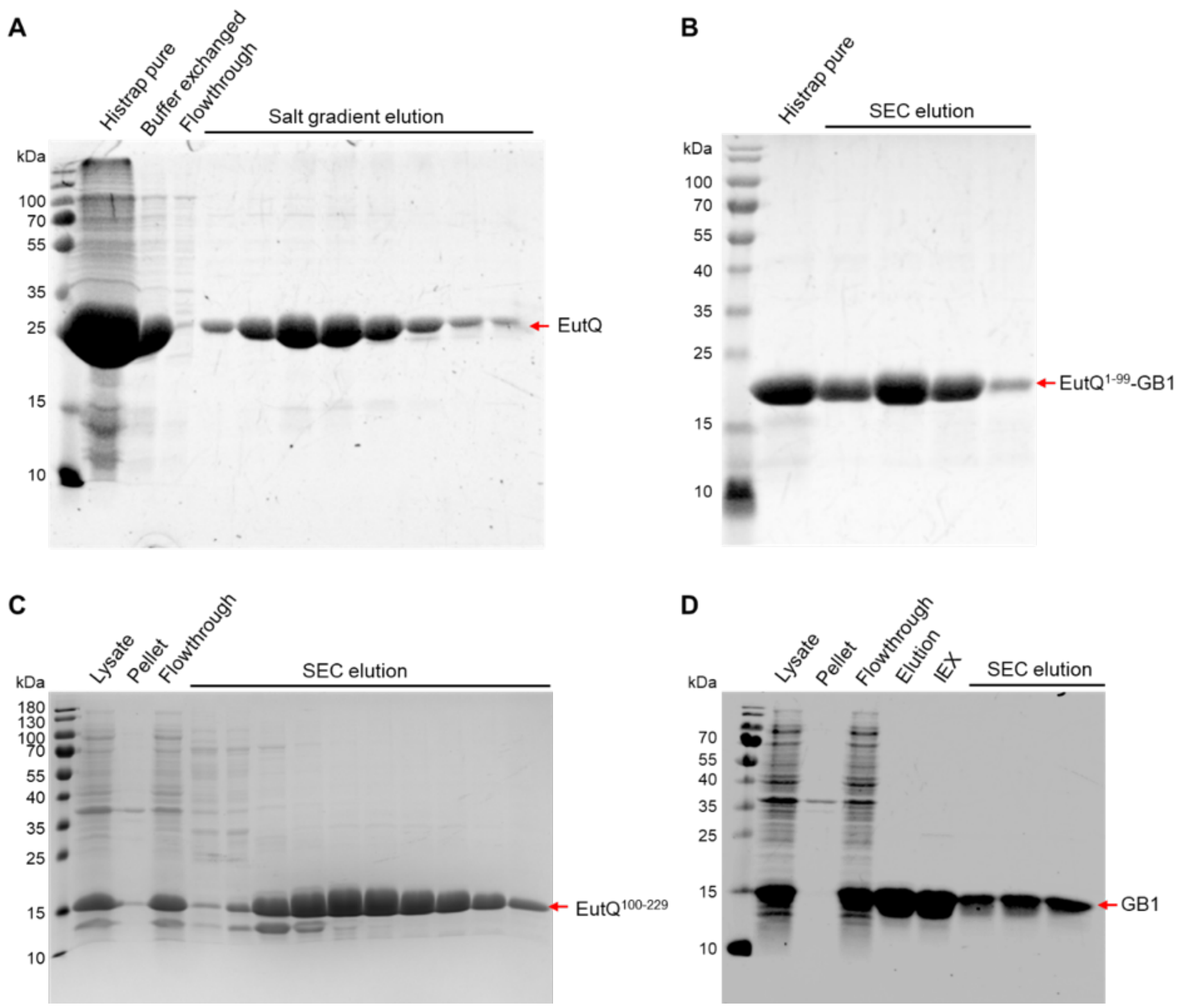


**Figure S17. Purification of EutQ (A), EutQ$^{1\text{-}99}$-GB1 (B), EutQ$^{100\text{-}229}$ (C), and GB1(D).** IEX: Ion exchange chromatography; SEC: Size exclusion chromatography.

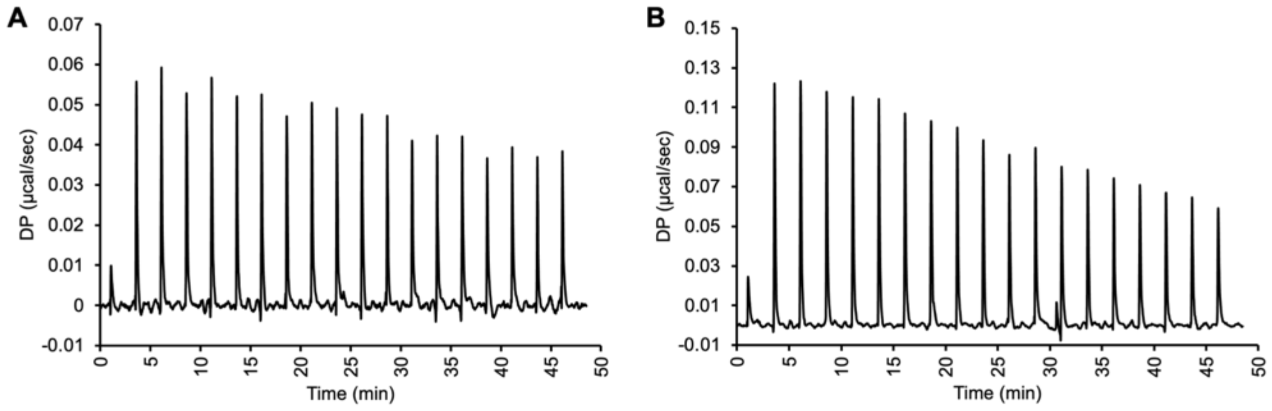


**Figure S18**. **ITC isotherm for controls. A**, EutC$^{1\text{-}20}$ to GB1 titration. **B**, EutC$^{1\text{-}20}$ to buffer titration. There was no binding in both experimental procedures, hence, no calculated best fit curve.

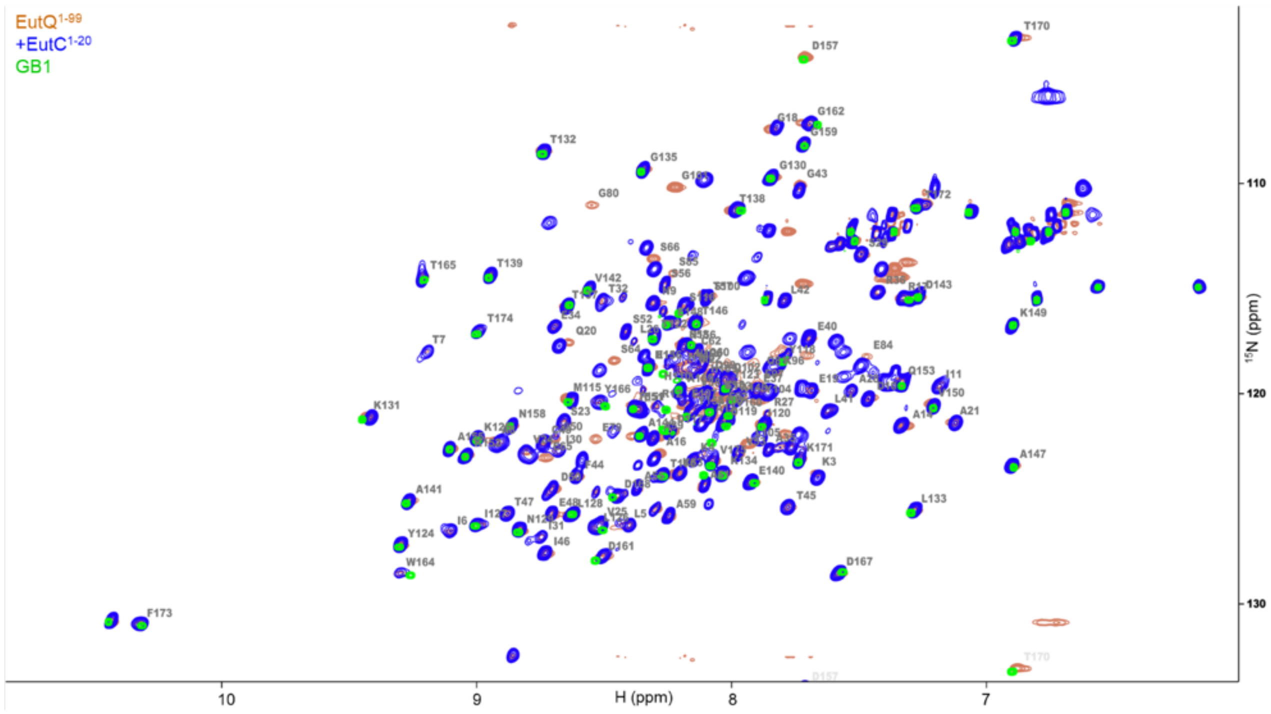


**Figure S19. Backbone assigned 2D $^{15}N$-$^{1}H$ HSQC spectrum of uniformly $^{15}N$-labeled EutQ$^{1\text{-}99}$-GB1 (red) (100 mM, pH 6.5, 298 K) overlaid with (i) $^{15}N$-$^{1}H$ HSQC spectrum of $^{15}N$ uniformly GB1 (green), and (ii) $^{15}N$ EutQ$^{1\text{-}99}$-GB1 in the presence of 500 mM unlabeled EutC$^{1\text{-}20}$ (blue).** The chemical shift perturbations that were observed in EutQ$^{1\text{-}99}$-GB1 without equivalent shifts on GB1 in the presence of ligand are consistent with specific interactions occurring between the EutQ$^{1\text{-}99}$ alone and EutC$^{1\text{-}20}$. Please note that the numbers of residues shown in this figure are shifted 1 residue due to an additional residue at position 1 (start codon ATG for methionine).

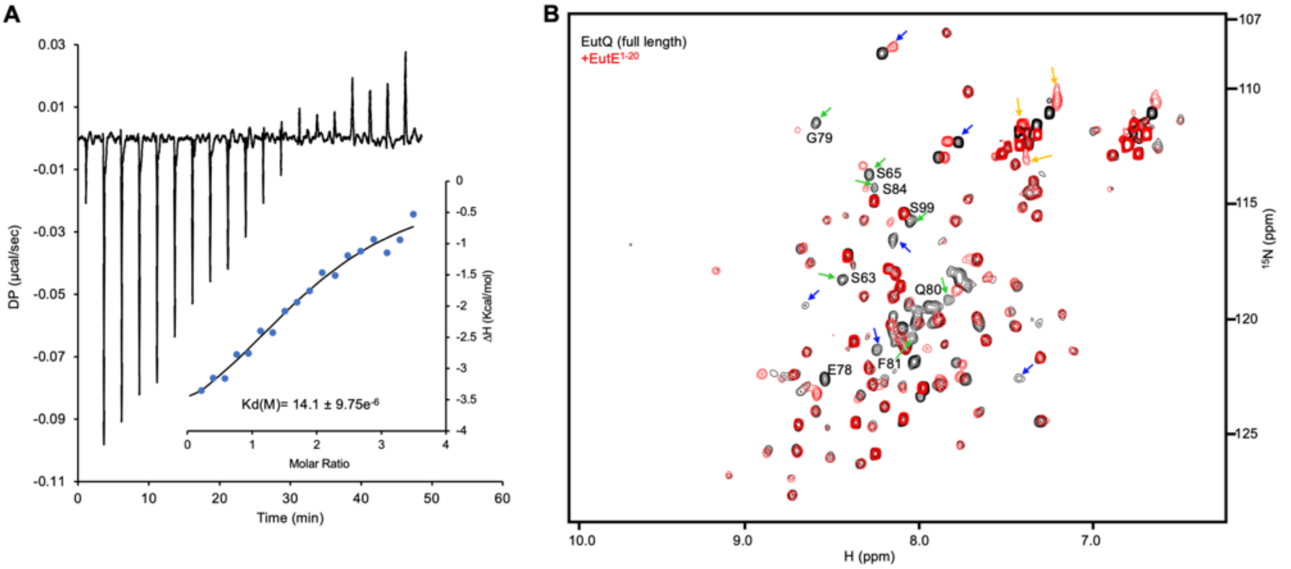


**Figure S20**. **EutE$^{1-20}$ interacts with EutQ$^{1-99}$. A**, Fitted isotherm of EutE$^{1-20}$ binding with EutQ (N ~ 2 with the peptide binding to each monomeric subunit in the EutQ dimer; $K_d$ = 14.1 μM; ΔG = -6.62 Kcal mol$^{-1}$; ΔH = -4.66 Kcal mol$^{-1}$; TΔS = -1.96 Kcal mol$^{-1}$). **B**, 2D $^{15}$N-$^{1}$H HSQC spectrum (298K) of uniformly $^{15}$N-labeled EutQ (black) overlaid with both $^{15}$N-labeled EutQ mixed with five-fold excess unlabeled EutE$^{1-20}$ (red). Assigned peaks are those derived from the EutQ N-terminal domain residues, which show chemical shift perturbations in the presence of EutE$^{1-20}$. Assigned peaks (63, 65, 78-81, 84, 99) chemical shift perturbations are highlighted with green arrows; unassigned peaks perturbations with blue arrows; Gln and Asn amide side chains with yellow arrows. Severe linewidth broadening precluded the assignments of many resonances in H3 and H4 regions.

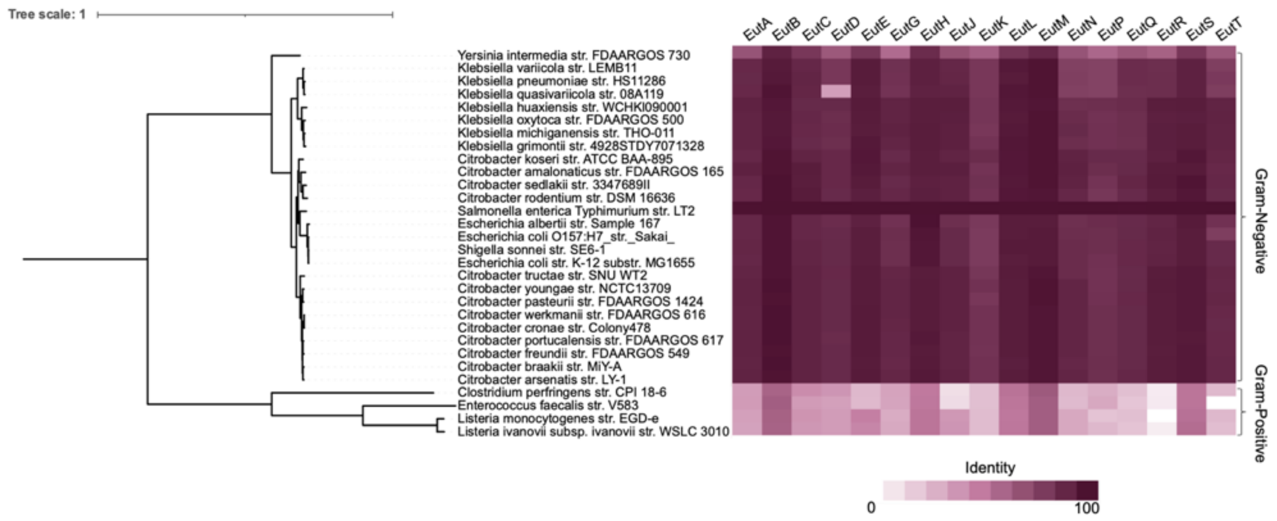


**Figure S21. Similarities among Eut proteins from different bacterial species at the amino acid level.** The phylogenetic tree was made from an alignment of 30 representative isolates containing the Eut BMCs (see Supplemental File 1). The tree was rooted on the middle point of Gram-negative and Gram-positive bacteria because they are too far away. The heatmap shows the similarity of all 17 Eut protein sequences of the 30 genomes comparing to *S.* Typhimurium LT2. Eut protein sequences are highly conserved among Gram-negative bacteria, whereas the similarity of Eut proteins between *S*. Typhimurium LT2 and Gram-positive bacteria is low.

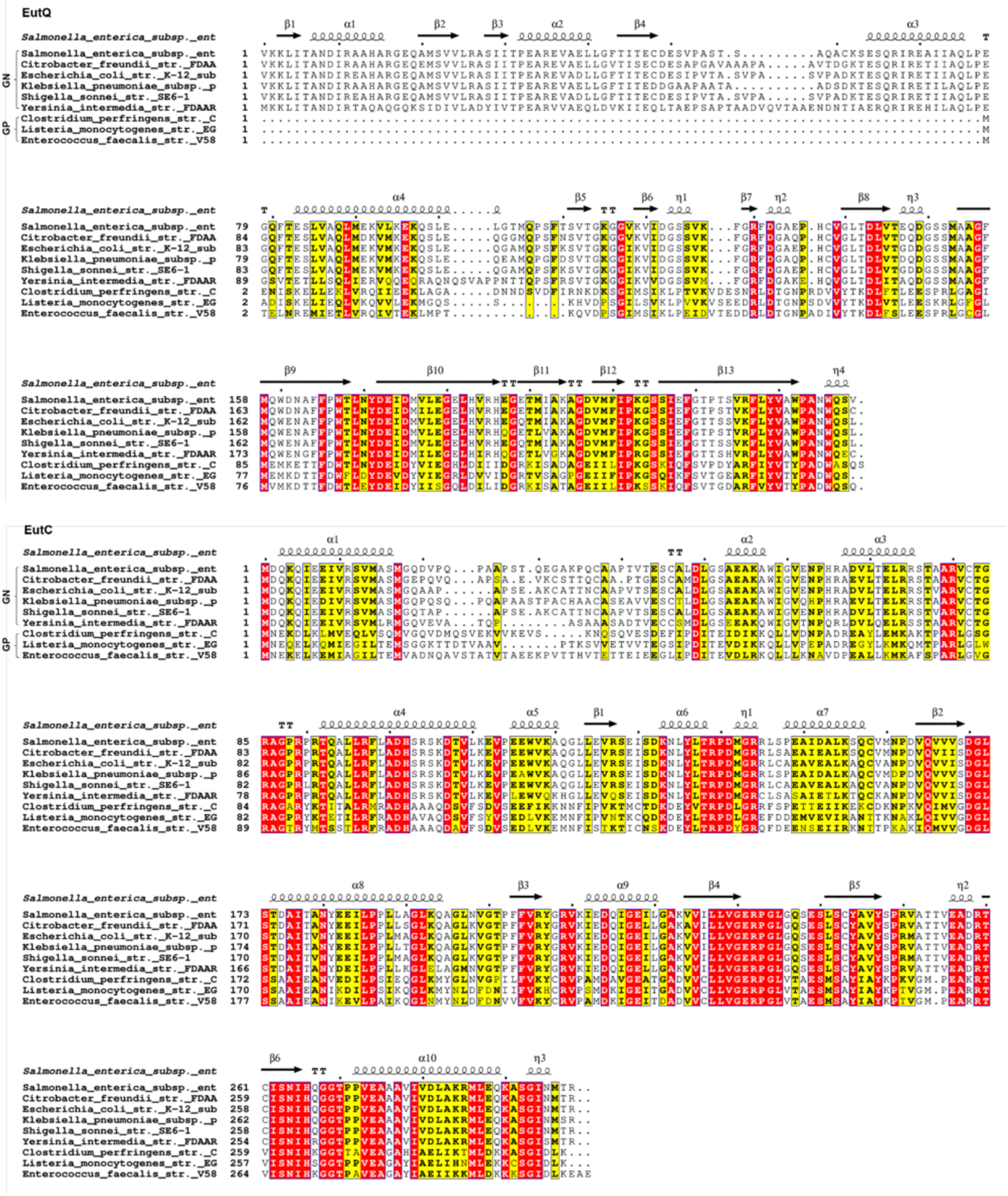
EutQ
EutC
GN
GP

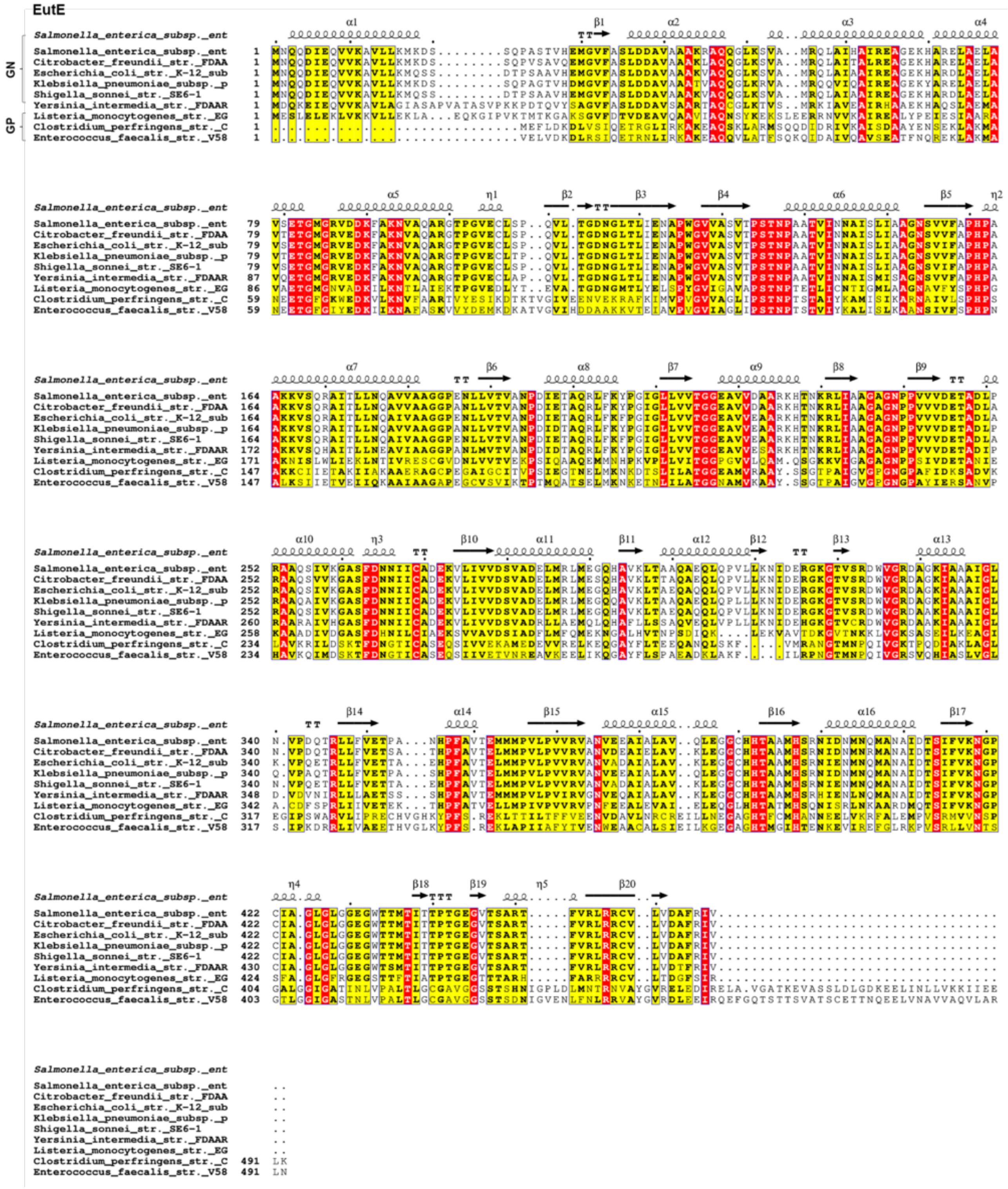


**Figure S22. Sequence alignment of EutQ, EutC and EutE from different bacterial species.** Eut protein structures are highly conserved among Gram-negative (GN) bacteria, whereas the similarity of Eut proteins between S. Typhimurium LT2 and Gram-positive (GP) bacteria is low. Red boxes indicate completely conserved amino acid residues and similar residues are written with black bold characters and boxed in yellow.

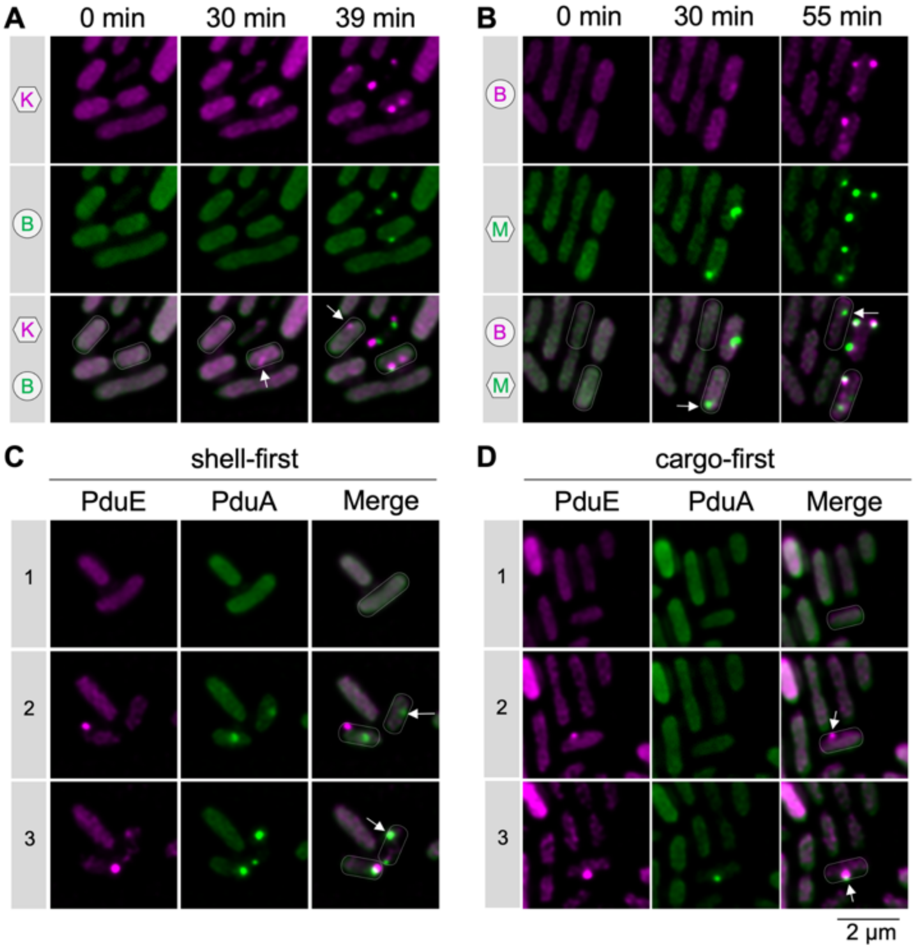


**Figure S23. Different assembly pathways of Eut BMC and Pdu BMC. A** and **B**, Aggregation of the shell (EutK-mCherry or EutM-sfGFP) and cargo (EutB-sfGFP or EutB-mCherry) of Eut BMC in the WT strain following induction with EA and $B_{12}$. **C** and **D**, Aggregation of the shell (PduA-sfGFP) and cargo (PduE-mCherry) of Pdu BMC in the WT strain following induction with 1,2-propanediol.

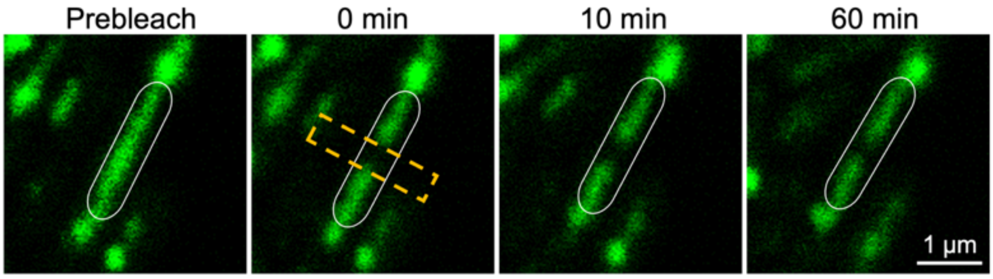


**Figure S24. Representative FRAP images of EutM-sfGFP (shell) at various time lapses.** The yellow rectangular boxes indicate the bleaching area.

**Table S1. Strains and plasmids used in this study.** Relevant antibiotic resistances are indicated by $^R$: Ap, ampicillin; Km, kanamycin; Cm, chloramphenicol; Gm, gentamicin; Tc, tetracycline.

| Strains/Plasmids | Description | Reference/Origin |
|---|---|---|
| LT2 | LT2, WT | (McClelland et al., 2001) |
| LT2-Δ*eutS* | Δ*eutS* | This study |
| LT2-Δ*eutN* | Δ*eutN* | This study |
| LT2-Δ*eutL* | Δ*eutL* | This study |
| LT2-Δ*eutK* | Δ*eutK* | This study |
| LT2-Δ*eutM* | Δ*eutM* | This study |
| LT2-Δ*eutQ* | Δ*eutQ* | This study |
| LT2-Δ$eutQ^{9\text{-}99}$ | Δ$eutQ^{9\text{-}99}$ | This study |
| LT2-Δ$eutQ^{100\text{-}229}$ | Δ$eutQ^{100\text{-}229}$ | This study |
| LT2-Δ$eutQ^{9\text{-}41}$ | Δ$eutQ^{9\text{-}41}$ | This study |
| LT2-Δ$eutQ^{42\text{-}61}$ | Δ$eutQ^{42\text{-}61}$ | This study |
| LT2-Δ$eutQ^{62\text{-}99}$ | Δ$eutQ^{62\text{-}99}$ | This study |
| LT2-Δ$eutQ^{9\text{-}21}$ | Δ$eutQ^{9\text{-}21}$ | This study |
| LT2-Δ$eutQ^{22\text{-}41}$ | Δ$eutQ^{22\text{-}41}$ | This study |
| LT2-Δ$eutQ^{62\text{-}79}$ | Δ$eutQ^{62\text{-}79}$ | This study |
| LT2-Δ$eutQ^{80\text{-}99}$ | Δ$eutQ^{80\text{-}99}$ | This study |
| LT2-Δ$eutE^{2\text{-}21}$ | Δ$eutE^{2\text{-}21}$ | This study |
| LT2-Δ$eutC^{2\text{-}19}$ | Δ$eutC^{2\text{-}19}$ | This study |
| LT2-Δ*eutP* | Δ*eutP* | This study |
| LT2-Δ*eutPQ* | Δ*eutPQ* | This study |
| ***E. coli* derivatives:** | | |
| *E. coli* S17-1 *λpir* | *pro thi hsdR recA* chromosome::RP4-2 Tc::Mu Km::Tn7/*λpir*; $Tp^R$, $Sm^R$ | (Simon et al., 1983) |
| **Plasmids:** | | |
| pEMG | Suicide plasmid; $Km^R$ | (Martínez-García and de Lorenzo, 2011) |
| pSW-2 | Plasmid for m-toluate–inducible expression of the I-SceI enzyme; $Gm^R$ | (Martínez-García and de Lorenzo, 2011) |
| pBAD/*Myc*-His | Vector for dose-dependent expression of recombinant proteins; $Ap^R$ | Invitrogen |
| pXG10-SF | Plasmid served as the backbone for complementation experiments; pSC101* origin of replication; $P_{LtetO\text{-}1}$ promoter; $Cm^R$ | (Corcoran et al., 2012) |
| pXG10-*eutK* | Plasmid for expression of EutK; $Cm^R$ | This study |
| pXG10-*eutL* | Plasmid for expression of EutL; $Cm^R$ | This study |
| pXG10-*eutQ* | Plasmid for expression of EutQ; $Cm^R$ | This study |
| pBAD-*MB* | *eutM::mCherry-eutB::sfGFP* cloned into pBAD/*Myc*-His at NcoI and HindIII sites; $Ap^R$ | This study |
| pBAD-*KB* | *eutK::mCherry-eutB::sfGFP* cloned into pBAD/*Myc*-His at NcoI and HindIII sites; $Ap^R$ | This study |
| pBAD-*LB* | *eutL::mCherry-eutB::sfGFP* cloned into pBAD/*Myc*-His at NcoI and HindIII sites; $Ap^R$ | This study |
| pBAD-*EB* | *eutE::mCherry-eutB::sfGFP* cloned into pBAD/*Myc*-His at NcoI and HindIII sites; $Ap^R$ | This study |

| | | |
|---|---|---|
| pBAD-*GB* | *eutG::mCherry-eutB::sfGFP* cloned into pBAD/*Myc*-His at NcoI and HindIII sites; $Ap^R$ | This study |
| pBAD-*PB* | *eutP::mCherry-eutB::sfGFP* cloned into pBAD/*Myc*-His at NcoI and HindIII sites; $Ap^R$ | This study |
| pBAD-*JB* | *eutJ::mCherry-eutB::sfGFP* cloned into pBAD/*Myc*-His at NcoI and HindIII sites; $Ap^R$ | This study |
| pBAD-*RB* | *eutR::mCherry-eutB::sfGFP* cloned into pBAD/*Myc*-His at NcoI and HindIII sites; $Ap^R$ | This study |
| pBAD-*QB* | *eutQ::mCherry-eutB::sfGFP* cloned into pBAD/*Myc*-His at NcoI and HindIII sites; $Ap^R$ | This study |
| pBAD-$Q^{\Delta 9\text{-}99}B$ | $eutQ^{\Delta 9\text{-}99}$*::mCherry-eutB::sfGFP* cloned into pBAD/*Myc*-His at NcoI and HindIII sites; $Ap^R$ | This study |
| pBAD-$Q^{\Delta 9\text{-}41}B$ | $eutQ^{\Delta 9\text{-}41}$*::mCherry-eutB::sfGFP* cloned into pBAD/*Myc*-His at NcoI and HindIII sites; $Ap^R$ | This study |
| pBAD-$Q^{\Delta 42\text{-}61}B$ | $eutQ^{\Delta 42\text{-}61}$*::mCherry-eutB::sfGFP* cloned into pBAD/*Myc*-His at NcoI and HindIII sites; $Ap^R$ | This study |
| pBAD-$Q^{\Delta 62\text{-}99}B$ | $eutQ^{\Delta 62\text{-}99}$*::mCherry-eutB::sfGFP* cloned into pBAD/*Myc*-His at NcoI and HindIII sites; $Ap^R$ | This study |
| pBAD-$E^{\Delta 2\text{-}21}B$ | $eutE^{\Delta 2\text{-}21}$*::mCherry-eutB::sfGFP* cloned into pBAD/*Myc*-His at NcoI and HindIII sites; $Ap^R$ | This study |
| pBAD-$C^{\Delta 2\text{-}19}B$ | $eutC^{\Delta 2\text{-}19}$*::mCherry-eutB::sfGFP* cloned into pBAD/*Myc*-His at NcoI and HindIII sites; $Ap^R$ | This study |
| pBAD-*BM* | *eutB::mCherry-eutM::sfGFP* cloned into pBAD/*Myc*-His at NcoI and HindIII sites; $Ap^R$ | This study |
| pBAD-*KM* | *eutK::mCherry-eutM::sfGFP* cloned into pBAD/*Myc*-His at NcoI and HindIII sites; $Ap^R$ | This study |
| pBAD-*LM* | *eutL::mCherry-eutM::sfGFP* cloned into pBAD/*Myc*-His at NcoI and HindIII sites; $Ap^R$ | This study |
| pBAD-*QM* | *eutQ::mCherry-eutM::sfGFP* cloned into pBAD/*Myc*-His at NcoI and HindIII sites; $Ap^R$ | This study |
| pBAD-$Q^{\Delta 62\text{-}99}M$ | $eutQ^{\Delta 62\text{-}99}$*::mCherry-eutM::sfGFP* cloned into pBAD/*Myc*-His at NcoI and HindIII sites; $Ap^R$ | This study |
| pBAD-eutBC-sfGFP | *eutBC::sfGFP* cloned into pBAD/Myc-His at NcoI and HindIII sites; $Ap^R$ | This study |
| pET14b-EutQ | *eutQ* cloned into pET14b backbone; $Ap^R$ | This study |
| pETM11-$EutQ^{1\text{-}99}$-GB1 | $eutQ^{1\text{-}99}$*-GB1* cloned into pETM11 backbone; $Km^R$ | This study |
| pETM11-$EutQ^{100\text{-}229}$ | $eutQ^{100\text{-}229}$ cloned into pETM11 backbone; $Km^R$ | This study |
| pETM11-GB1 | *GB1* cloned into pETM11 backbone; $Km^R$ | This study |

**Table S2. ssDNA Oligonucleotides used in this study.**

| Primers | Sequence (5'→3') |
|---|---|
| eutS-del-F1 | AGGGATAACAGGGTAATCTGAATTCAGGACTACATAAAGCACCAG |
| eutS-del-R1 | CCGCCACCAATTAACTTTTTGTCATTGCTCACCGTCTCCAC |
| eutS-del-F2 | GTGGAGACGGTGAGCAATGACAAAAAGTTAATTGGTGGCG |
| eutS-del-R2 | CCTGCAGGTCGACTCTAGAGGATCGTAATCGTGAAGCCCAGTAG |
| eutM-del-F1 | AGGGATAACAGGGTAATCTGAATTCGGATATCGCTCTCGCCA |
| eutM-del-R1 | CTTATCCGGCCTACCATCACATCGTGTTTTCCTCTCATTAAA |
| eutM-del-F2 | TTTAATGAGAGGAAAACACGATGTGATGGTAGGCCGGATAAGA |
| eutM-del-R2 | CCTGCAGGTCGACTCTAGAGGATCCTGCGTCATCCAGGGAG |
| eutN-del-F1 | AGGGATAACAGGGTAATCTGAATTCGCTAATTCAGGGGCTTGC |
| eutN-del-R1 | CATGATGTTCAATCCTATTTATGCATGGATTGCCCGTTGGGTC |
| eutN-del-F2 | GACCCAACGGGCAATCCATGCATAAATAGGATTGAACATCATG |
| eutN-del-R2 | CCTGCAGGTCGACTCTAGAGGATCAGAGTGATTGCCCGCTGA |
| eutL-del-F1 | AGGGATAACAGGGTAATCTGAATTGTGCGTTCGGAGATCAGC |
| eutL-del-R1 | CAGCCTCCGTTACGCACGCATGATGTCTCCTTAACGGG |
| eutL-del-F2 | CCCGTTAAGGAGACATCATGCGTGCGTAACGGAGGCTG |
| eutL-del-R2 | CCTGCAGGTCGACTCTAGAGGATCCGTCTTCGGGTAACGCTTC |
| eutK-del-F1 | AGGGATAACAGGGTAATCTGAATTCCTATATTGCCGCTGACGAAG |
| eutK-del-R1 | CCCGGCCCTCCGACAGATTACATTGGCAGCCTCCGTTA |
| eutK-del-F2 | TAACGGAGGCTGCCAATGTAATCTGTCGGAGGGCCGGG |
| eutK-del-R2 | CCTGCAGGTCGACTCTAGAGGATCCATATGCAGCACCCTTTCC |
| eutC2-19-del-F1 | AGGGATAACAGGGTAATCTGAATTCAAGGGTCTGCGCTCTCC |
| eutC2-19-del-R1 | CGGGCTGCGGTACGTCCATGGTGTTATCCCCGCG |
| eutC2-19-del-F2 | CGCGGGGATAACACCATGGACGTACCGCAGCCC |
| eutC2-19-del-R2 | CCTGCAGGTCGACTCTAGAGGATCGGCGTGCCGACGTTCAG |
| eutE2-21-del-F1 | AGGGATAACAGGGTAATCTGAATT GCAAAGCCGCAACCGATG |
| eutE2-21-del-R1 | GAACAGTACTGGCCGGCTGCATGATGTTCAATCCTATTTATGG |
| eutE2-21-del-F2 | CCATAAATAGGATTGAACATCATGCAGCCGGCCAGTACTGTTC |
| eutE2-21-del-R2 | CCTGCAGGTCGACTCTAGAGGATC GTATGTTTACGCGCCGCG |
| eutQ-del-F1 | AGGGATAACAGGGTAATCTGAATTGCGCTATTACAGACGGTCAG |
| eutQ-del-R1 | CGGATTGCCAGTTTGCAGGCTAGTTAGCTGTGATAAGTTTTTTCACC |
| eutQ-del-F2 | GGTGAAAAAACTTATCACAGCTAACTAGCCTGCAAACTGGCAATCCG |
| eutQ-del-R2 | CCTGCAGGTCGACTCTAGAGGATCCTCATCCTCGTTCAGTCC |
| eutL-del-9-99-R1 | CATCGTCCCCAGTTCCAGGTTAGCTGTGATAAGTTTTTTCACC |
| eutL-del-9-99-F2 | GGTGAAAAAACTTATCACAGCTAACCTGGAACTGGGGACGATG |
| eutL-del-100-229-R1 | CGGATTGCCAGTTTGCAGGCTACGACTGCTTTTCCTTCAGC |
| eutL-del-100-229-F2 | GCTGAAGGAAAAGCAGTCGTAGCCTGCAAACTGGCAATCCG |
| eutL-del-9-21-R1 | GCGCAGCACTACCGAGTTAGCTGTGATAAGTTTTTTCACC |
| eutL-del-9-21-F2 | GGTGAAAAAACTTATCACAGCTAACTCGGTAGTGCTGCGC |
| eutL-del-9-41-R1 | CATTCGGTAATCGTGAAGCCGTTAGCTGTGATAAGTTTTTTCACC |
| eutL-del-9-41-F2 | GGTGAAAAAACTTATCACAGCTAAC GGCTTCACGATTACCGAATG |
| eutL-del-22-41-R1 | CATTCGGTAATCGTGAAGCCCATCGCCTGTTCGCCGCGGG |
| eutL-del-22-41-F2 | CGGCGAACAGGCGATGGGCTTCACGATTACCGAATG |
| eutL-del-42-61-R1 | TGCGCTGGCTTTCGCTTTTCAGTAGTTCCGCCACTTCG |
| eutL-del-42-61-F2 | CGAAGTGGCGGAACTACTG AAAAGCGAAAGCCAGCGCA |
| eutL-del-62-79-R1 | CACCAGGCTTTCAGTAAACTGACACGCCTGCGCCGATG |
| eutL-del-62-79-F2 | TCGGCGCAGGCGTGT CAGTTTACTGAAAGCCTGGTG |
| eutL-del-80-99-R1 | CATCGTCCCCAGTTCCAGCCCTTCCGGGAGCTG |

| | |
|---|---|
| eutL-del-80-99-F2 | CAGCTCCCGGAAGGG CTGGAACTGGGGACGATG |
| eutL-del-62-99-R1 | CATCGTCCCCAGTTCCAGACACGCCTGCGCCGATG |
| eutL-del-62-99-F2 | CATCGGCGCAGGCGTGT CTGGAACTGGGGACGATG |
| eutS-up | GCACAAATCGAGTGATACGC |
| eutS-down | CGCGTGGACATAAATCAGCG |
| eutM-up | GATGATTTTCCCGTCGCTGG |
| eutM-down | TACTGTCGATAGCGACGGC |
| eutN-up | CATTGGCGAGTTGGTCTCC |
| eutN-down | GAATGGCATGAATGGCAAGC |
| eutL-up | TCGTCATCCTGCTGGTAGG |
| eutL-down | CTTTGCGACTAATCACACGGC |
| eutK-up | GAAGCGACATTCGGCATTG |
| eutK-down | GATCGTAGATCTGCTGCCAG |
| eutE2-21-up | ATCGATGCCCAGGGCAAC |
| eutE2-21-down | GCGGAGATAAACACTCGACG |
| eutC2-19-up | GCTGTAACTACATCATGGGGA |
| eutC2-19-down | CCAGGAAGCGCAACAGC |
| eutQ-up | GGAACCCGTAAACGACATATC |
| eutQ-down | GTTAAGCCATGTACCGGC |
| pBAD-eutMB-F1 | GGGCTAACAGGAGGAATTAACCATGGAAGCATTAGG |
| pBAD-eutMB-R1 | CGTCCTCGAAGTTCATCACGCGCTCCCACTTGAAG |
| pBAD-eutMB-F2 | CTTCAAGTGGGAGCGCGTGATGAACTTCGAGGACG |
| pBAD-eutMB-R2 | ATTCCTAATGCTTCCATGGTTAATTCCTCCTGTTAGCCCCTACTTG |
| pBAD-eutMB-F3 | AACCATGGAAGCATTAGGAATGATTGAAACCCGGGGCCTGGTTGCGCTGATT GAGGCCT |
| pBAD-eutMB-R3 | TTTGCTCATGAATTCGCCAGAACCAGCAGCGGAGCCAGCGGATCCAATGTTG CTGTCGCC |
| pBAD-eutMB-F4 | CTGGCGAATTCATGAGCAAAG |
| pBAD-eutMB-R4 | CTGAGATGAGTTTTTGTTCTACGTA |
| pBAD-mCherry-eutB/M-sfGFP-F1 | CTGGCTCCGCTGCTGGTTCTGGCGAATTCGTGAGCAAGGGCGAGGAG |
| pBAD-mCherry-eutB/M-sfGFP-R1 | GAGATGAGTTTTTGTTCTACGTATTATTTGTAGAGCTCATCCATGCCA |
| pBAD-eutE-F1 | GGGCTAACAGGAGGAATTAACCATGAATCAACAGGATATTGAACAGG |
| pBAD-eutE-R1 | AGAACCAGCAGCGGAGCCAGCGGATCCTACAATGCGAAACGCATCCA |
| pBAD-eutE-KO2-21-F1 | GGGCTAACAGGAGGAATTAACCATGCAGCCGGCCAGTACTGTTC |
| pBAD-eutP-F1 | GGGCTAACAGGAGGAATTAACCATGAAACGTATTGCTTTTGTCG |
| pBAD-eutP-R1 | AGAACCAGCAGCGGAGCCAGCGGATCCGCTGTGATAAGTTTTTTCACCTG |
| pBAD-eutQ-F1 | GGGCTAACAGGAGGAATTAACCATGAAAAAACTTATCACAGCTAAC |
| pBAD-eutQ-R1 | AGAACCAGCAGCGGAGCCAGCGGATCCTACGGATTGCCAGTTTGC |
| pBAD-eutJ-F1 | GGGCTAACAGGAGGAATTAACCATGGCGCACGACGAACA |
| pBAD-eutJ-R1 | AGAACCAGCAGCGGAGCCAGCGGATCCGCTTGCATAGAGTCCCTCC |
| pBAD-eutR-F1 | GGGCTAACAGGAGGAATTAACCATGAAAAAGACCCGTACAGC |
| pBAD-eutR-R1 | AGAACCAGCAGCGGAGCCAGCGGATCCAGCCCATTGCCGCATCC |
| pBAD-eutK-F1 | GGGCTAACAGGAGGAATTAACCATGATCAATGCCCTGGGA |
| pBAD-eutK-R1 | AGAACCAGCAGCGGAGCCAGCGGATCCATTTTTGATGCGATAGCGACTA |
| pBAD-eutL-F1 | GGGCTAACAGGAGGAATTAACCATGCCTGCATTAGATTTAATTCG |
| pBAD-eutL-R1 | AGAACCAGCAGCGGAGCCAGCGGATCCCGCACGCTGGACAGG |
| pBAD-eutG-F1 | GGGCTAACAGGAGGAATTAACCATGCAAGCTGAACTACAGAC |
| pBAD-eutG-R1 | AGAACCAGCAGCGGAGCCAGCGGATCCCCCGGCAGCCGCGTAC |
| pBAD-eutC-KO-2-19-F1 | GGGCTAACAGGAGGAATTAACCATGGACGTACCGCAGCCCGCC |
| pBAD-eutC-R1 | AGAACCAGCAGCGGAGCCAGCGGATCCACGGGTCATGTTGATGCCG |

| | |
|---|---|
| pBAD-BC-sfGFP-F1 | GGGCTAACAGGAGGAATTAACCATGAAACTAAAGACCACATTGTTCGG |
| pBAD-BC-sfGFP-R1 | GGAGCCAGCGGATCCACGGGTCATGTTGATGC |
| pBAD-BC-sfGFP-F2 | CCCGTGGATCCGCTGGCTCCG |
| pBAD-BC-sfGFP-R2 | GAGATGAGTTTTTGTTCTACGTAAGCTT |
| pXG10-eutL-F | GTGATAGAGATACTGAGCACATGCATTCCGGCATCAACATGACCCG |
| pXG10-eutL-R | CTTTCGTTTTATTTGATGCCTCTAGATTACGCACGCTGGACAGGGTTA |
| pXG10-eutQ-F | GTGATAGAGATACTGAGCACATGCATTGGCGGCTCTCAGTGAACA |
| pXG10-eutQ-R | CTTTCGTTTTATTTGATGCCTCTAGATCATACGGATTGCCAGTTTGC |
| pXG10-eutK-F | GTGATAGAGATACTGAGCACATGCATAACCCTGTCCAGCGTGC |
| pXG10-eutK-R | CTTTCGTTTTATTTGATGCCTCTAGATTAATTTTTGATGCGATAGCGACTAC |
| pXG10-eutC-F | GTGATAGAGATACTGAGCACATGCATTCACTGTTCTTCTGATGACG |
| pXG10-eutC-R | CTTTCGTTTTATTTGATGCCTCTAGATTAACGGGTCATGTTGATGC |
| EutQ_For | TTTTCAGGGCAAGAGGCAGGTGAAAAAACTTATCACAGC |
| EutQ_Rev | AGCCGGATCCTCATACGGATTGCCAGTTTGCAGG |
| pET14b_EutQ_For | ATCCGTATGAGGATCCGGCTGCTAACAAAGCC |
| pET14b_EutQ_Rev | CCTGCCTCTTGCCCTGAAAATACAGGTTTTCCATATG |
| EutQ_C_Term_For | TTTTCAGGGCCTGGAACTGGGGACGATGCAG |
| EutQ_C_Term_Rev | CGGATCCCTATCATACGGATTGCCAGTTTGCAGG |
| pETM11_EutQ_C_Term_For | ATCCGTATGATAGGGATCCGAATTCGAGCTCCGTC |
| pETM11_EutQ_C_Term_Rev | CCAGTTCCAGGCCCTGAAAATAAAGATTCTCAGTAGTGGGGATG |
| EutQ_N_Term_For | ATATACCATGGTGAAAAAACTTATCACAGCTAACGATATTCGTGCGG |
| EutQ_N_Term_Rev | TCAGGGCTCACGACTGCTTTTCCTTCAGCACTTTTTC |
| GB1_His_Tev_For | AAAGCAGTCGTGAGCCCTGAAAATACAGGTTTTCCATCACCATCACCATCACCCCATGAG |
| GB1_His_Tev_Rev | ATCCCTATTATTCGGTCACGGTAAAGGTTTTGG |
| pETM11_EutQ_N_Term_For | CGTGACCGAATAATAGGGATCCGAATTCGAGCTCC |
| pETM11_EutQ_N_Term_Rev | GTTTTTTCACCATGGTATATCTCCTTCTTAAAGTTAAACAAAATTATTTCTAGAGGG |
